# A transient postnatal quiescent period precedes emergence of mature cortical dynamics

**DOI:** 10.1101/2021.02.17.430487

**Authors:** Soledad Domínguez, Liang Ma, Han Yu, Gabrielle Pouchelon, Christian Mayer, George D. Spyropoulos, Claudia Cea, György Buzsáki, Gord Fishell, Dion Khodagholy, Jennifer N. Gelinas

**Author notes:** Corresponding authors: Dion Khodagholy, Jennifer Gelinas. These authors contributed equally to this manuscript (Soledad Domínguez, Liang Ma, Han Yu).

## Abstract

Mature neural networks synchronize and integrate spatiotemporal activity patterns to support cognition. Emergence of these activity patterns and functions is believed to be developmentally regulated, but the postnatal time course for neural networks to perform complex computations remains unknown. We investigate the progression of large-scale synaptic and cellular activity patterns across development using high spatiotemporal resolution *in vivo* electrophysiology in immature mice. We reveal that mature cortical processes emerge rapidly and simultaneously after a discrete but volatile transition period at the beginning of the second postnatal week of rodent development. The transition is characterized by relative neural quiescence, after which spatially distributed, temporally precise, and internally organized activity occurs. We demonstrate a similar developmental trajectory in humans, suggesting an evolutionarily conserved mechanism to transition network operation. We hypothesize that this transient quiescent period is a requisite for the subsequent emergence of coordinated cortical networks.

## Introduction

Multiple cognitive functions emerge rapidly during early development. Neural networks enable precise spatiotemporal coordination of synaptic and cellular activity in mature brain functions (*1–4*). How immature neural networks develop into their mature form needed for the complex computations underlying cognition remains poorly understood. The first organized and predominant pattern of neural activity that appears in cortical circuits across species is a spindle-like oscillation (10-20 Hz) that occurs intermittently on a background of relative neural inactivity. Known as spindle bursts in rodents and delta brushes in humans, this immature network activity is commonly triggered by peripheral stimuli (*5–7*). They demarcate cortical columns or pre-columns, and have been linked to neuronal survival (*8*), establishing sensory ensembles, and critical period plasticity (*7, 9–11*). Spindle bursts are characteristic of the first postnatal week of rodent development, and delta brushes disappear shortly after term in human neonates (*12*), emphasizing their transient role in network maturation. In contrast, mature cortex exhibits perpetual, complex patterns of neural activity that appear and interact across a wide range of frequencies (*13–15*). The organization of this activity creates precise spatiotemporal windows for neural synchronization, enabling plasticity processes and generation of neural sequences (*16–18*). Such network properties facilitate information processing, and resultant activity patterns have been causally linked to cognitive processes from stimulus perception to learning and memory (*1–4*). Understanding when and how these properties develop in the immature brain is critical given that delay or failure to express mature brain activity is a strong risk factor for subsequent impaired cognition (*7, 19–21*). We hypothesized that emergence of these advanced neural network properties could be heralded by the disappearance of immature spindle activity patterns. During this developmental epoch, cortical microcircuits are still undergoing dramatic changes in anatomical connectivity (*7*) and functional connectivity between and within cortical layers is initiated (*22*) in part through an abrupt increase in synaptogenesis in superficial layers (*23*). Robust GABA-mediated fast inhibition also appears (*24, 25*) as inhibitory network influences of interneurons increase (*26–28*). These local circuit changes are paralleled with the gradual ingrowth of subcortical neurotransmitters that assist in setting brain state changes. Together, such modifications could prime a shift in network operating mode, setting the stage for re-emergence of spindle band activity in the form of thalamocortical sleep spindles, a characteristic pattern of the offline state of non-rapid eye movement (NREM) sleep that facilitates memory consolidation(*2, 29*).

To investigate this hypothesis, we examined neural network dynamics across early rodent and human development. We targeted epochs spanning disappearance of immature activity and emergence of mature sleep patterns in each species. For rodents, we developed conformable, minimally invasive, high resolution electrocorticography arrays coupled with high-density implantable probes and recorded spontaneous *in vivo* electrophysiologic patterns from unanesthetized mice across the first two postnatal weeks. This approach enabled simultaneous acquisition of large-scale synaptic and cellular activity without damaging fragile cortical circuits. We found that the transition between immature and mature network dynamics was characterized by unexpectedly decreased coordinated cellular and synaptic activity at the beginning of the second postnatal week. After this timepoint, precise neuronal synchronization and oscillatory coupling in space and time robustly emerged. Analysis of continuous electroencephalography (EEG) recordings from human subjects, 36 to 69 weeks post-gestation, revealed a similar developmental trajectory characterized by a transient decrease in neural activity prior to onset of spatiotemporal oscillatory coupling. Therefore, a shift from local, loosely correlated, prominently sensory-driven patterns to internally organized, spatially distributed and temporally precise activity is preceded by a transient quiescent period. These findings suggest an evolutionarily conserved mechanism to developmentally regulate functional network capacity.

## Results

To identify and characterize emergence of advanced neural network properties in the developing brain, we acquired high spatiotemporal resolution electrophysiological recordings from somatosensory cortex of unanesthetized mouse pups aged postnatal day (P) 5 to 14 (n = 108 pups), an epoch that spans the transition from immature to mature activity patterns during sleep. We used minimally invasive surface electrocorticography arrays (NeuroGrids, n = 70 pups) to permit a spatially extensive survey of cortex. These customized NeuroGrids (*30*) consisted of 119 electrodes regularly spaced on a diagonal square centered lattice embedded in 4 µm thick parylene C to conform to the cortical surface (**Figure 1A**). Recordings were made following recovery from surgery to eliminate any influence of anesthesia. Mice at these ages enter into a cyclical pattern of sleep and wakefulness as assayed by peripheral indicators, such as muscular tone, movements, and heart rate (**Supplementary Figure 1**). We ensured electrodes used for analysis were located in somatosensory cortex by post-mortem histology. Following NeuroGrid recordings, the location of the array was marked on the surface of the brain using a biocompatible fluorescent material (*31*) and tissue was harvested for immuno-histochemistry. vGlut2 staining identified the location of primary somatosensory and visual cortices relative to NeuroGrid placement on flattened axial slices (**Figure 1B, upper**). This histology-based electrode grouping corresponded to spatial localization of electrophysiological activity patterns, identifying electrodes recording from somatosensory cortex (**Figure 1B, lower; Supplementary Figure 2**). To additionally capture neural spiking patterns across cortical layers, we stereotactically implanted silicon probes (linear electrode arrangement) in somatosensory cortex of a separate cohort (n = 38 pups). Immunohistochemistry of coronal slices from these brains verified probe placement within somatosensory cortex, permitted allocation of recording electrodes to cortical layers (**Figure 1C, upper**), and demonstrated transcortical neural spiking patterns (**Figure 1C, lower, Supplementary Figure 3**).

**Figure 1:**
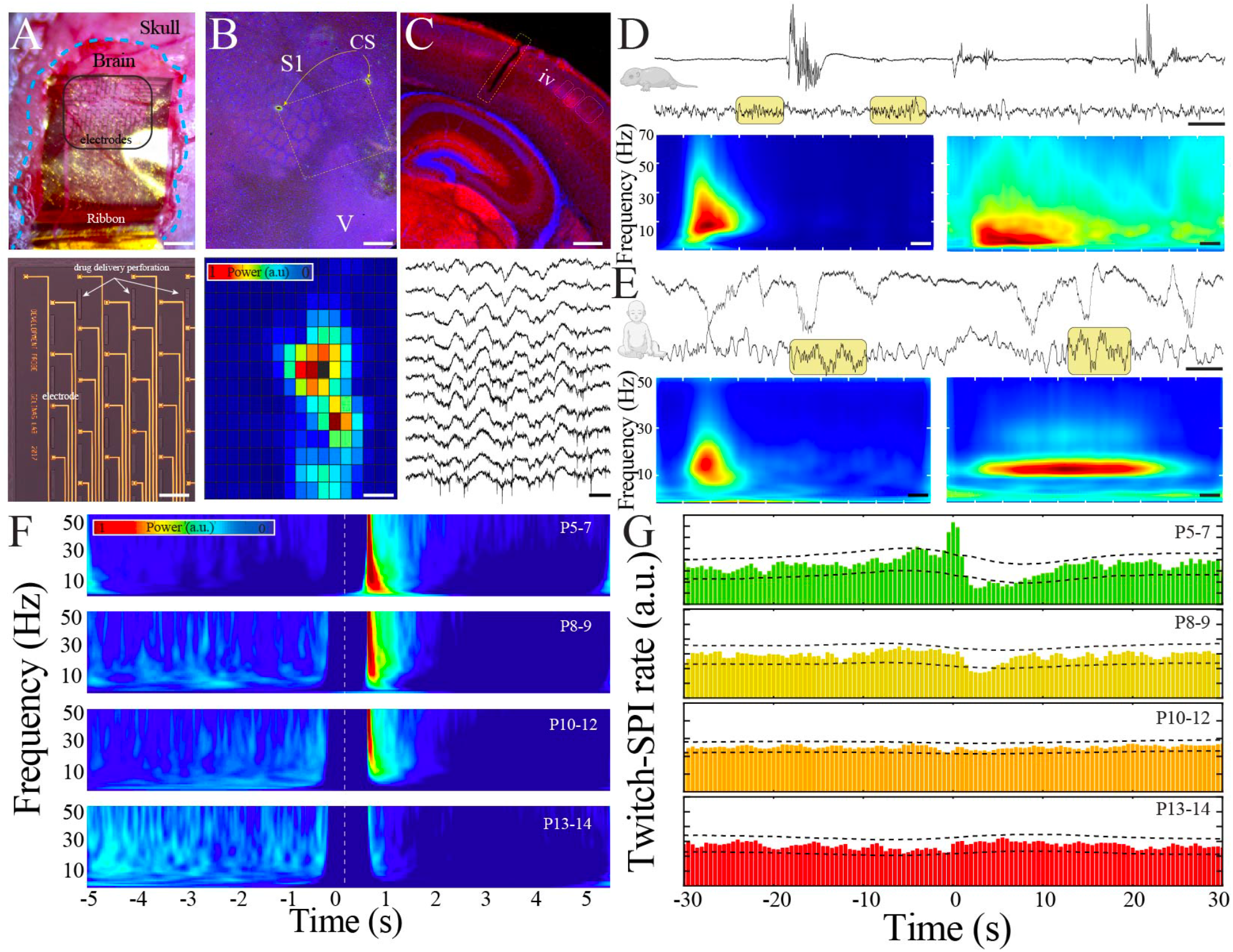
Large-scale, high spatiotemporal resolution recording of neural activity across development in rodents demonstrates conserved features with human EEG. A) Optical micrograph of NeuroGrid conforming to the surface of a P7 mouse pup (upper; scale bar 200 µm). Optical micrograph revealing arrangement of electrodes and perforations in section of NeuroGrid (lower; scale bar 50 µm). B) Fluorescence microscopy of flattened brain slice from P14 mouse pup demonstrating location of NeuroGrid during recording (yellow dashed lines). vGlut2 immunohistochemistry enables identification of visual and somatosensory cortices. S1 = primary somatosensory; V = visual; CS = chitosan marking corners of NeuroGrid (upper; scale bar 200 µm). Spatial distribution of spindle band power across a NeuroGrid array in a P7 mouse pup. Warmer colors signify higher power (lower; scale bar 500 ms). C) Fluorescence microscopy of coronal brain slice from P7 mouse pup demonstrating location of implantable silicon probe (yellow dashed lines). vGlut2 immunohistochemistry enables identification of barrels in granular layer of cortex (IV) (upper; scale bar 200 µm). Raw traces across a sample of silicon probe electrodes demonstrating localized neural spiking activity in a P13 mouse pup (lower; scale bar 200 ms). D) Sample raw traces from P5 (upper) and P14 (lower) mouse pups showing shift from discontinuous to continuous activity. Yellow boxes highlight sleep spindles; scale bar 1 s. Spectrograms trigger-averaged on spindle band oscillations in P5 (left) and P14 (right) mouse pups (n = 50 oscillations from each sample session; scale bar = 200 ms). E) Sample raw traces from 37 week post-gestational age (upper) and 58 week post-gestational age (lower) human subjects showing shift from discontinuous to continuous activity. Yellow boxes highlight sleep spindles; scale bar 1 s. Spectrograms trigger averaged on spindle-band oscillations in 37 week post-gestational age (left) and 58 week post-gestational age (right) subjects (n = 50 oscillations from each sample session; scale bar = 1 s). F) Twitch-triggered spectrograms (twitch center at time = 0, white dashed line) show a progressive decrease in movement evoked oscillatory activity across development (n = 6716 twitches from n = 52 pups). Note power temporally coincident with twitch is set to zero to avoid any potential artifactual contamination. G) Cross-correlation of twitch and detected spindle band oscillation (SPI) decreases across development. Dashed lines are 95% confidence intervals; time 0 = twitch (n = 6716 twitches from n = 52 pups).

We expected that physiologically significant features of neural network maturation would be conserved across species. Thus, in parallel, we obtained continuous EEG recordings from normal human subjects 36-69 weeks post-gestation (1 day-7 months after birth), an epoch that similarly spans the transition from immature to mature activity patterns of human sleep and can be mapped to corresponding timepoints of rodent brain development (*32–34*). We first identified epochs of artifact-free behavioral immobility consistent with sleep in the pups (**Supplementary Figure 4**). In line with electrophysiological patterns that have been observed in preterm human neonates, sleep in the youngest pups (P5-7) was characterized by irregular, discontinuous bursts of neural activity (**Figure 1D-E, upper traces**). The predominant activity was organized into spindle bursts, oscillations characteristic of developing rodent sensory cortex (**Figure 1D, left spectrogram**). These spindle bursts share morphological and frequency characteristics with delta brushes, an oscillatory pattern of the human brain that develops by 26 weeks post-gestation and disappears around 40-44 weeks (*35*) (**Figure 1E, left spectrogram)**. More mature pups exhibited nearly continuous patterns during sleep, an absence of spindle bursts, and the appearance of sleep spindles, which are well characterized oscillations of NREM sleep (**Figure 1D, lower trace, right spectrogram**). Human sleep is similarly continuous and sleep spindles appear during NREM between 1-2 months after term in infants (**Figure 1E, lower trace, right spectrogram**). This mature sleep in rodents and humans is predominantly governed by internally generated dynamics (*36*). In contrast, self-generated or evoked movements during sleep are closely associated with LFP activity in somatosensory cortex of immature rodents and humans (*7, 37*). We confirmed that the capacity for muscular twitches to evoke changes in LFP power (**Figure 1F;** n = 6716 twitches from n = 52 pups, sampled randomly to ensure uniform group size; Kruskal-Wallis ANOVA, Chi Square = 12.93, p = 0.0048) or detectable spindle band oscillations (**Figure 1G;** cross-correlation of twitches and spindle band oscillations exceeded 95% confidence interval, black dashed lines, only in pups aged P5-7) decreased as the pups matured. These marked similarities in the electrophysiological features of neural activity during sleep across development suggest the possibility of evolutionarily conserved mechanisms of neural network maturation.

To investigate how this maturation occurs, we examined the macrostructure of sleep LFP with fine temporal precision over early pup development (P5-14). Mature sleep can be classified into distinct stages (NREM and REM) based on electrophysiological criteria, but these indicators do not reliably differentiate the immature analogues of these states (quiet and active sleep, respectively) until after P11(*20*). Therefore, we focused on periods of behavioral and EMG quiescence that lasted for longer than 10 seconds (**Supplementary Figure 4**), with the goal of preferentially analyzing the transition of quiet sleep (which contains spindle bursts) into NREM sleep (which contains sleep spindles). Unexpectedly, recordings across development revealed an epoch of relative neural quiescence at the beginning of the second postnatal week (**Figure 2A**) that corresponded to decreased spectral power and absence of prominent activity at physiologic frequencies (**Figure 2B**). We hypothesized that this quiescence could be related to a transient decrease in the continuity of the neural signal or the power of the activity that was present. Continuity of neural activity was quantified by determining the duration of neural activity above the wideband noise floor. Although continuity increased markedly from the youngest to most mature animals, we found a decrease in the duration of continuous neural activity that corresponded to the identified period of spectral quiescence (**Figure 2C**). When we extracted the average wideband power of the continuous activity that was present at each developmental day, there was also a transient nadir at this timepoint (**Figure 2D**). Because we aimed to characterize the trajectory of the electrophysiological features over time without making a priori assumptions about the nature of the temporal relationship or the rate of change, we employed a modeling approach that considered age as a continuous variable. This approach avoided arbitrary grouping of data from different aged pups without sacrificing statistical power, and enabled characterization of the developmental profile beyond testing at individual timepoints. Data were fit with linear and polynomial regression models, and model performance was evaluated using leave-one-out cross-validation (LOOCV) to avoid over-fitting (**Supplementary Figure 5**). Polynomial regression provided the best fit for continuity and power data as quantified by the mean squared error (MSE; **Supplementary Figure 5B**). In addition, linear regression was insufficient to model these parameters because linear fit residuals systematically deviated from zero and consistently overestimated observed values at the beginning of the second postnatal week (**Supplementary Figure 5C**). The best fit models exhibited local minima, and to estimate the age at which these nadirs occurred, we employed a bootstrapping method. Bootstrapping data within each developmental day localized probability of nadir in continuity and power at the beginning of the second postnatal week with likelihood far exceeding the distribution obtained by bootstrapping across days (**Figure 2C-D inset, Supplementary Figure 5D-F, Supplementary Figure 6A**). Therefore, sleep LFP macrostructure expresses a nonlinear pattern of development that is characterized by a discrete period of relative neural quiescence occurring at the beginning of the second postnatal week.

**Figure 2:**
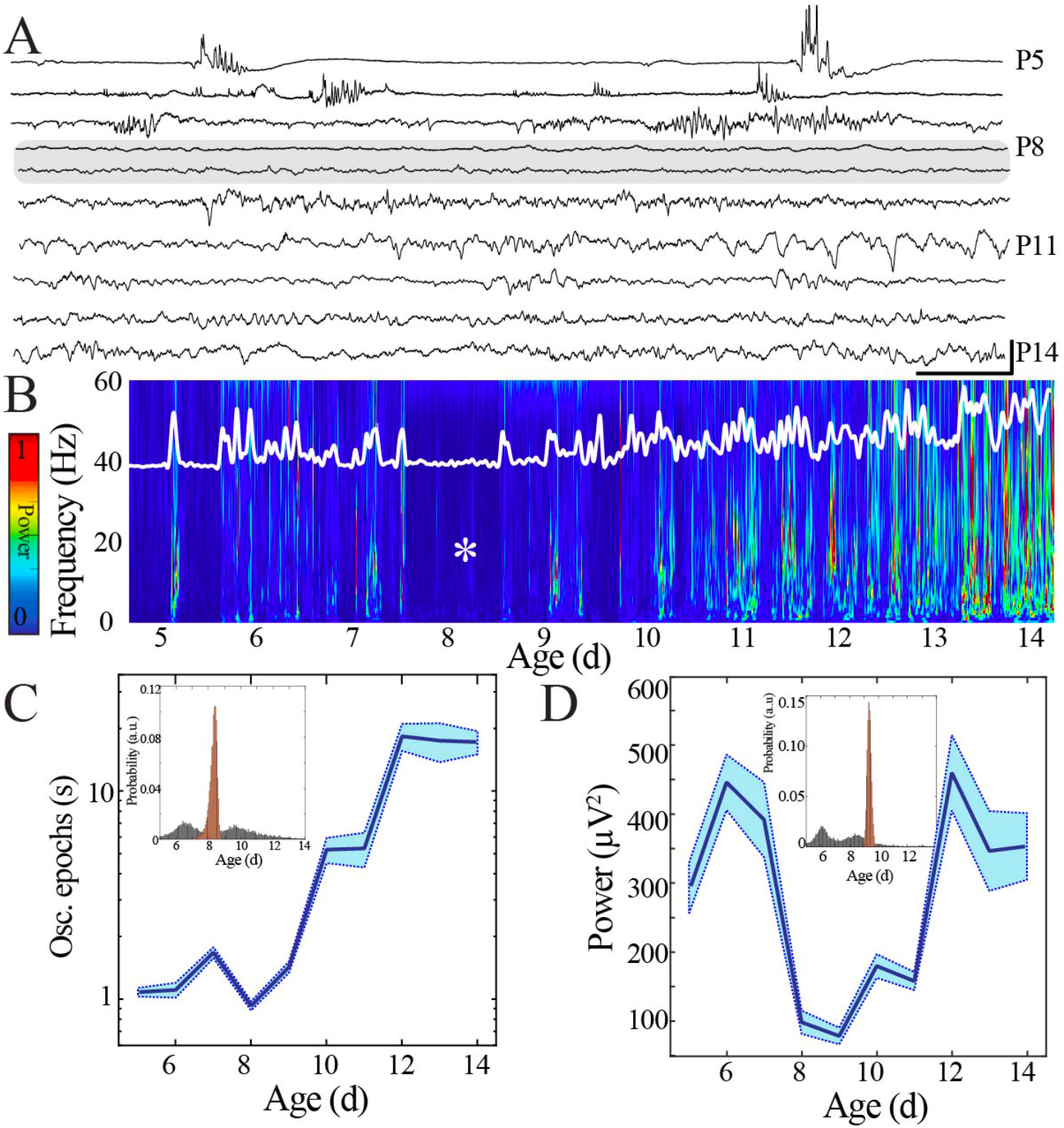
Duration and power of cortical oscillatory activity transiently decreases at the beginning of the second postnatal week. A) Sample raw NeuroGrid traces from P5-14 mouse pups demonstrating changing characteristics of oscillatory patterns across development with relative paucity of activity at P8 and P9 (shaded gray box). Scale bar 1 s, 250 µV. B) Compilation of individual spectrograms (60 s duration) from a sample session per day of age demonstrates a transient reduction in LFP power at the beginning of the second postnatal week. The white superimposed trace represents the overall instantaneous power of neural activity. C) Duration of oscillatory epochs increases non-linearly across development (blue line = mean; shaded blue areas ± SE) with a local minimum at the beginning of the second postnatal week (n = 70 pups). Inset shows probability of a local minimum located at each age for datapoints resampled with replacement across days (grey) compared to datapoints resampled with replacement within days (orange), confirming existence of local minimum between P8-9. D) Wideband (WB) power changes non-linearly across development (blue line = mean; shaded blue areas ± SE), with a local minimum at the beginning of the second postnatal week (n = 70 pups). Inset shows probability of a local minimum located at each age for datapoints resampled with replacement across days (grey) compared to datapoints resampled with replacement within days (orange), confirming existence of local minimum between P8-9.

We then sought to understand the functional relevance of this quiescent transition period to network operations. Key LFP properties (spatial extent, waveform, and power) can provide an index of the underlying neural population activity and its computational purpose - the distance over which neurons can be concurrently recruited, as well as the strength and temporal patterning of this activity (*38–41*). We detected spindle band oscillations, which were present in pups of all ages, to facilitate comparisons. We quantified spatial extent using the two-dimensional organization of the NeuroGrid to determine the proportion of electrodes expressing co-occurring spindle band oscillations with a reference electrode located in primary somatosensory cortex (**Figure 3A**) (*42*). Waveform was characterized using an asymmetry index (**Figure 3B**) and power was extracted from the filtered Hilbert envelope (**Figure 3C**). Spindle band oscillations in the most immature pups were high power and had a substantially asymmetric waveform, but were expressed over a restricted cortical area (∼480 µm diameter). In contrast, spindle band oscillations in the most mature pups were lower power and more symmetric, but could variably be detected over a larger spatial extent (∼1.4 mm diameter). Given these observed differences in oscillatory parameters, we asked to what extent they could be used to classify the developmental stage at which the oscillations occurred. A coarse tree model based on spatial extent and waveform asymmetry was effectively trained to classify individual spindle band oscillations as originating from immature (P5-7) or mature (P10-14) pups with 92.4% accuracy (**Figure 3D**), suggesting that these parameters capture defining features of the oscillations at the ends of the developmental spectrum. Spindle band oscillations that occurred during the transient quiescent period at the beginning of the second postnatal week were not consistently classified into either group, instead displaying a unique combination of oscillatory parameters: large spatial extent, low power, and intermediate waveform asymmetry (**Figure 3A-C**, orange bars). When examined in a combinatorial fashion, a significant inflection point in oscillation properties emerged between P8-9 (**Figure 3E**). Polynomial regression provided the best fit for these data (**Supplementary Figure 6B)**. Bootstrapping data within each developmental day confirmed localization of a nadir at the beginning of the second postnatal week with likelihood far exceeding the distribution obtained by bootstrapping across days (**Figure F**). These data indicate that oscillations are transformed at the beginning of the second postnatal week, potentially reflecting a transition in neural network function from consistently highly localized cortical processing to capacity for broader, tuned recruitment of neural activity.

**Figure 3:**
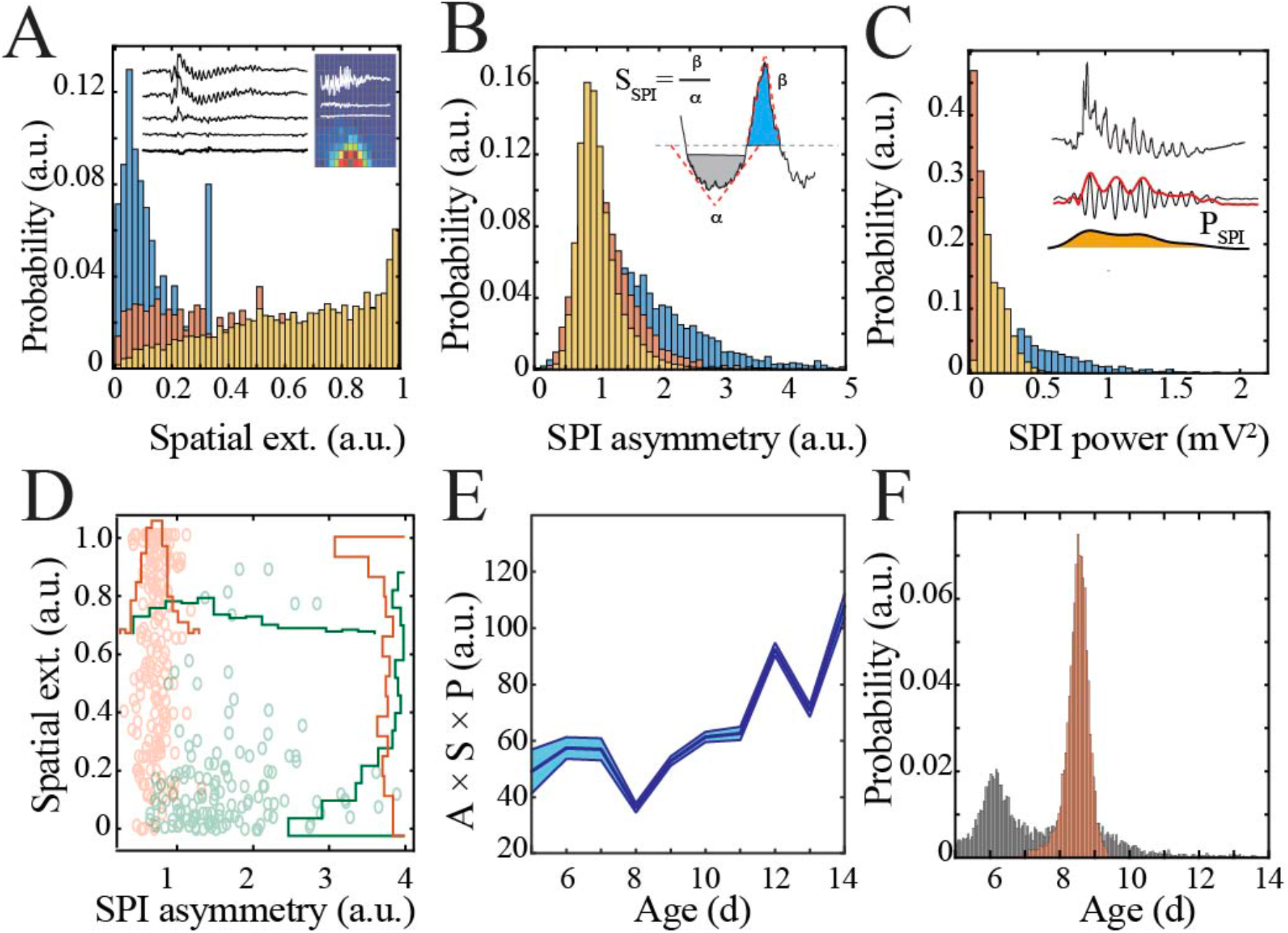
Key functional oscillatory properties transition at the beginning of the second postnatal week. A) Magnitude and variability of spindle band oscillation spatial extent varies across pup development (n = 44 pups, 42335 spindle band oscillations; Kolmogorov-Smirnov (KS) statistic P5 vs. P8 = 0.47, p < 0.001; P5 vs. P14 = 0.71, p < 0.001; P8 vs. P14 = 0.26, p < 0.001; all p-values adjusted for multiple comparisons using Bonferroni correction). Inset shows sample raw traces from spatially distributed NeuroGrid electrodes (left) and average spindle band power across NeuroGrid array (each square represents one NeuroGrid electrode; warmer traces indicate higher power) from a P5 pup. B) Waveform asymmetry of spindle band oscillations decreases across pup development (n = as in 3A); Kolmogorov-Smirnov (KS) statistic P5 vs. P8 = 0.37, p < 0.001; P5 vs. P14 = 0.52, p < 0.001; P8 vs. P14 = 0.20, p < 0.001; all p-values adjusted for multiple comparisons using Bonferroni correction). Inset shows the process of extracting the angles associated with peak and trough as well as formula for asymmetry index. C) Power of spindle band oscillations decreases across pup development (n = as in 3A); Kolmogorov-Smirnov (KS) statistic P5 vs. P8 = 0.69, p < 0.001; statistic P5 vs. P14 = 0.42, p < 0.001; P8 vs. P14 = 0.54, p < 0.001; all p-values adjusted for multiple comparisons using Bonferroni correction). Inset demonstrates process of extracting spindle band power, from raw trace to filtered trace and extraction of Hilbert power envelope for sample spindle band oscillation from P5 pup. D) Coarse tree model trained on spatial extent and waveform asymmetry of manually classified spindle band oscillations (n = 315 training set) from immature (P5-7) and mature (P10-14) pups has 92.4% accuracy in clustering oscillations into putative immature spindle bursts (green) and mature sleep spindles (orange). Superimposed line plots demonstrate the marginal probability for each event type. E) Combinatorial metric of spindle band waveform properties (A = asymmetry; S = spatial extent; P = power) increases non-linearly across development (black line = mean; shaded blue areas ± SE) with a local minimum at the beginning of the second postnatal week (n = 44 pups). F) Probability of a local minimum located at each age for data points resampled with replacement across days (grey) compared to datapoints resampled with replacement within days (orange), confirming existence of local minimum between P8-9.

Oscillations in the adult brain precisely organize neural activity across spatial and temporal scales. A key marker for this organization is cross frequency coupling, which is important for regulating strength of inputs from different afferent pathways (*43*) and directing flow of information processing (*44*). The marked shift in oscillatory properties that we observed at the beginning of the second postnatal week suggested that this organizational capacity may also be developmentally regulated. We first investigated the relationship of spindle band activity with other frequencies by detecting epochs of sustained increase in spindle band power and calculating measures of amplitude cross frequency coupling. Spindle band epochs in the most immature animals (P5-7) demonstrated high comodulation across a broad, non-specific range of frequencies, in contrast to frequency specific comodulation expressed in mature animals (P13-14; **Figure 4A**). Specific, discrete frequency bands that demonstrated significant temporal coupling with spindle band oscillations emerged only after P8-9. Discrete coupling with the gamma band (45-60 Hz) first occurred at P10-12, with an additional high gamma band (60-90 Hz) arising at P13-14 (**Figure 4B**; n = 1020 spindle band oscillations from 51 pups; area under curve (AUC) for 45-60 Hz Chi Square = 10.13, p = 0.02; AUC for 60-90 Hz Chi Square = 24.29, p < 0.001). Taken together, these results suggest that immature synaptic activity lacks the capacity to couple discrete frequency bands, whereas after the beginning of the second postnatal week such capacity emerges, potentially signaling a developmental shift in the function of oscillations.

**Figure 4:**
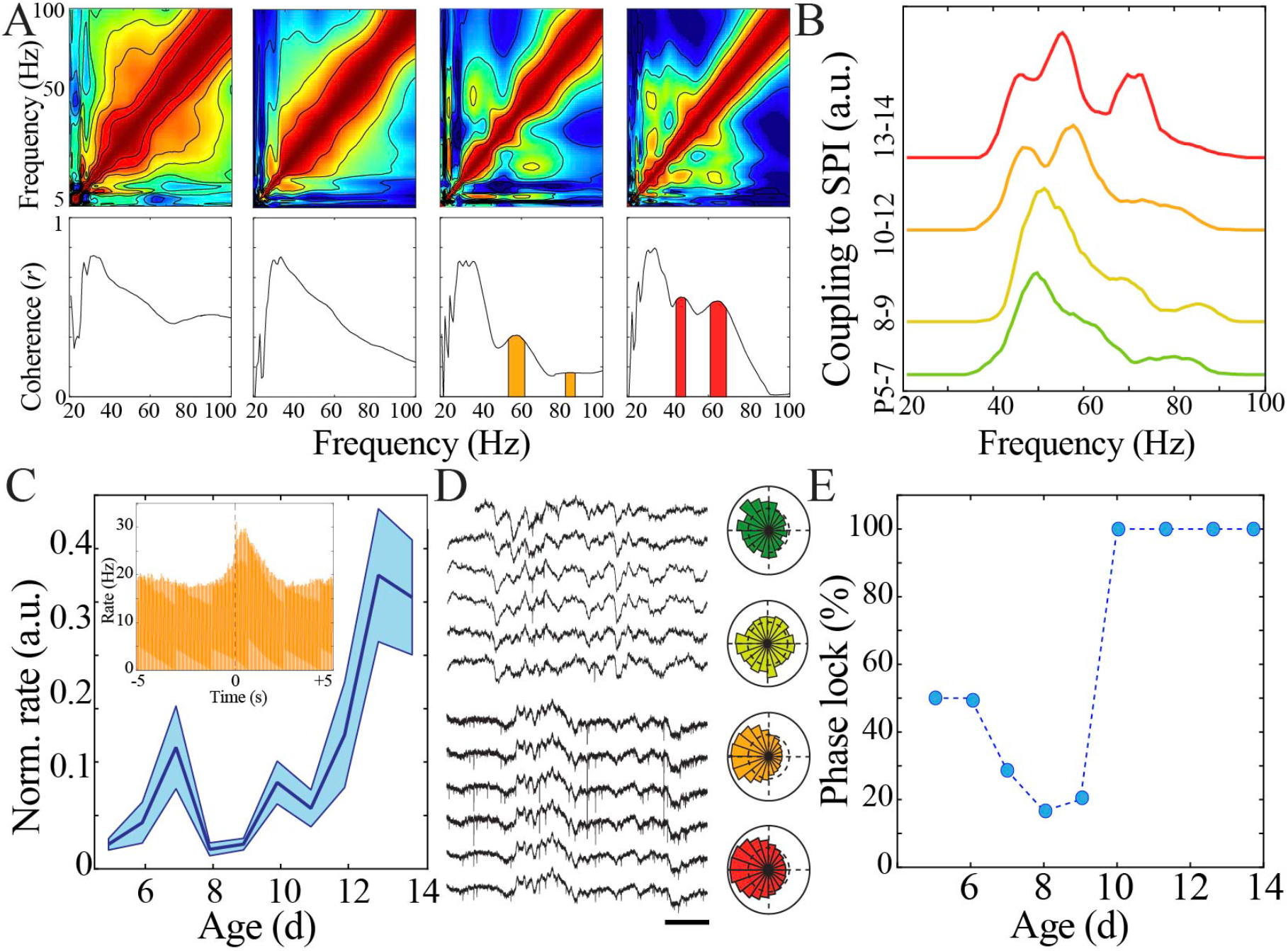
Increasing regularity and temporal precision of synaptic activity facilitates recruitment and entrainment of neural spiking across development. A) Frequency comodulation at the time of spindle band oscillations emerges across development. Comodulograms (upper) and quantification of significant peaks in frequency coherence to spindle band (lower). Scale bar 200 ms. B) Significant, discrete peaks of cross-frequency coupling with spindle band oscillations emerge starting at P10 (n = 51 pups). C) Rate of neural spiking during spindle band oscillation normalized to baseline neural spiking rate increases non-linearly across development (blue line = mean; shaded blue areas ± SE) with a local minimum at the beginning of the second postnatal week (n = 38 pups). Inset shows sample peristimulus time histogram of neural spiking during spindle band oscillation (starting at t = 0) for a P14 pup. D) Sample raw time traces of spindle-band oscillation and neural spiking from P8 (top) and P13 (bottom) pups. Prominent phase-locking of spikes to trough of spindle-band oscillation occurs in P13 pup only. Scale bar 500 ms. Sample polar plots show significant (kappa > 0.1 and Rayleigh p < 0.05) phase locking of neural firing to spindle band oscillations in P5 (green), P11 (orange), and P14 (red) pups, but not at P8 (yellow). E) Percentage of pups expressing significant (kappa > 0.1 and Rayleigh p < 0.05) phase-locking of neural spiking to spindle band oscillations across development (n = 38 pups).

Furthermore, oscillations facilitate neural communication by biasing spike occurrence to particular phases of the oscillatory waveform. We detected spiking activity recorded using implantable silicon probes from two non-contiguous sites (> 200 µm vertical separation) within the deep layers of cortex (IV-VI) per pup recording (n = 38 pups). Spindle band oscillations recruited neural spiking above baseline rates to a variable extent across development, with the lowest amount of recruitment occurring at the beginning of the second postnatal week (**Figure 4C**). This trajectory significantly deviated from linearity with a discrete nadir confirmed by bootstrapping during this epoch (**Supplementary Figure 7A**). P8 and P9 also exhibited the lowest proportion of recording sessions with significant phase-locking (kappa > 0.1 and alpha < 0.05), and consistency of phase-locking abruptly increased at P10 (**Figure 4D-E**; n = 38 pups implanted with silicon probes). These results reinforce the notion of enhanced information processing capacity in cortical networks after a rapid transitional period at the beginning of the second postnatal week.

Rate and synchrony of neural spiking strongly contribute to expression of oscillatory LFP patterns. We hypothesized that changes in neural spiking patterns could drive the shifts in neural network operation we observed. We first calculated rates of neural spiking only during epochs of above threshold LFP activity to eliminate effects related to LFP discontinuity. We found that neural spiking increased non-linearly over the course of development (**Figure 5A**), again with a significant nadir at the beginning of the second postnatal week as quantified using a bootstrapping method (**Supplementary Figure 7B**). These data suggest that lowered intracortical synaptic drive could contribute to the nadir in LFP patterns at the beginning of the second postnatal week. Consistent spiking within neuronal integration time is critical for plasticity processes (*45*). To estimate this property, we calculated the inter-spike-interval (ISI) between each unique action potential. ISI duration decreased non-linearly across development, with the longest ISI duration occurring in pups at the beginning of the second postnatal week (**Figure 5B, Supplementary Figure 7C**). Population neural spiking at fine time scales is thus both transiently decreased and desynchronized at this time, resulting in a brief period of relative quiescence prior to the progressive increase in spiking rate and synchrony that accompany subsequent maturation. During mature NREM sleep, there is alternation between periods of neural spiking and hyperpolarization which serves to synchronize neural populations and is implicated in memory consolidation (*46, 47*). We examined population spiking dynamics in the developing cortex by computing the autocorrelation of spiking activity, which permits quantification of neural synchrony over durations relevant to this slow oscillation (**Figure 5C**). Significant short time scale (< 50 ms) interactions (above the 95% confidence interval) robustly emerged only after P9 and were prominent in the most mature pups (**Figure 5D, left;** n = 38 pups, rank sum z = −4.2, p = 2.5×10^−5^). All age groups demonstrated significant suppression of spiking activity within the first 500 ms, becoming more marked as development progressed (**Figure 5D, right;** n = 38 pups, rank sum z = 3.0, p = 0.002). These data implicate that the transient quiescent period demarcates a transition in population spiking that initiates development of a cycle of longer scale network synchronization characteristic of the dynamics observed during mature sleep (*47*).

**Figure 5:**
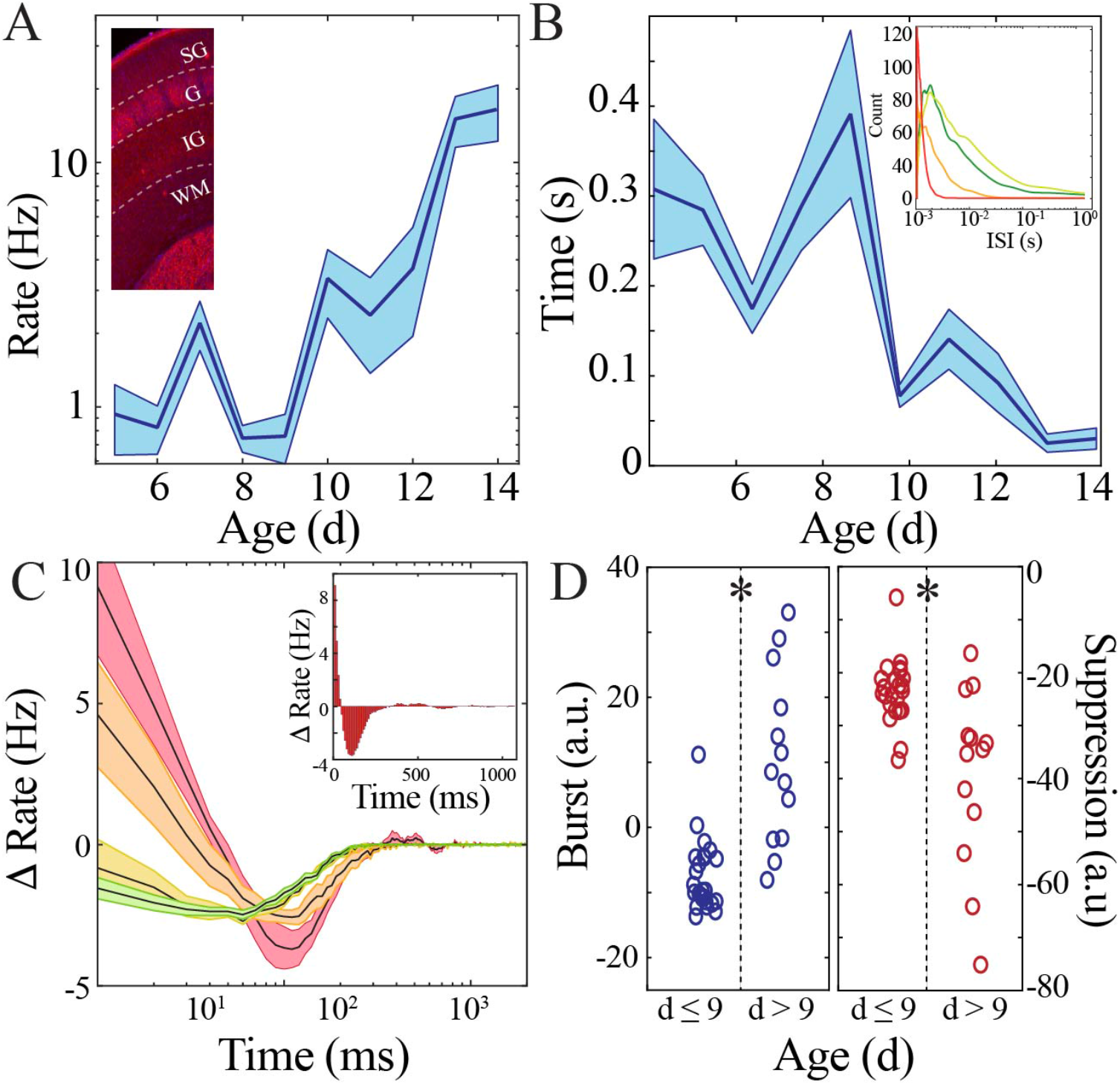
Rate and temporal precision of neural spiking increase after the beginning of the second postnatal week. A) Neural spiking rate (layers IV-VI) increases non-linearly across development (blue line = mean; shaded blue areas ± SE) with a local minimum at the beginning of the second postnatal week (n = 38 pups). Inset shows that vGlut2 immunohistochemistry facilitates identification of layers in barrel cortex (left; SG = supragranular; G = granular; IG = infragranular; WM = white matter). B) Inter-spike interval of spiking activity decreases non-linearly across development (blue line = mean; shaded blue areas ± SE) with a local maximum at the beginning of the second postnatal week (n = 38 pups). Inset shows histogram distribution of 10000 inter-spike intervals randomly selected per age group (red = P13-14, orange = P10-12, yellow = P8-9, green = P5-7). C) Average amount of significantly positively correlated (> 95% upper confidence interval, corresponding to a rate change > 0) or negatively correlated (< 95% lower confidence interval, corresponding to a rate change < 0) neural spiking as computed using spike autocorrelation (traces are mean ± SE). Sample autocorrelation of population neural spiking used for deriving the significant correlations in a P14 pup (inset). Significantly correlated neural spiking at fine-time scales (< 50 ms) emerged starting at P10 (n = 38 pups; P5-7 = green, P8-9 = yellow; P10-12 = orange; P13-14 = red). D) Average amount of significant positive and negative correlation (outside of 95% confidence intervals) that occurs within 50 ms (burst window; n = 38 pups, z = −4.2, p = 2.5e^-5^) and 50-500 ms (suppression window; n = 38 pups, z = 3.0, p = 0.002) as computed using spike autocorrelation for immature and mature pups.

Thus, multiple LFP and neural spiking measures followed a similar highly non-linear developmental trajectory marked by a key transition at the beginning of the second postnatal week in mouse pups. We next aimed to investigate whether this pattern of neural maturation was conserved across species. To accomplish this goal, we analyzed clinically acquired continuous EEG data from 55 human subjects ranging from 36 to 69 weeks post-gestation. EEG recordings were included for analysis only if they were reported as normal by the reading epileptologists and the subject had no known underlying neurologic or genetic condition. The majority of subjects received a discharge diagnosis of normal movements/behaviors (e.g. sleep myoclonus) or brief resolved unexplained events (**Supplementary Figure 8**). We analyzed epochs of clinically determined quiet/NREM sleep from each subject. Raw traces and normalized spectral analysis suggested an epoch of transient neural quiescence between 42 to 47 weeks (**Figure 6A-B**), akin to what we observed at the beginning of the second postnatal week in mice. To quantify these observations, we again used regression fitting to determine whether a nonlinear trajectory was present and bootstrapping of data within and between ages to identify any significant nadirs in neural activity patterns. Wideband power of oscillatory activity followed a highly non-linear trajectory, with a significant nadir between 42-47 weeks (**Figure 6C, Supplementary Figure 9A**). As expected from previous work (*12*), duration of oscillatory epochs generally increased over development, but we did observe a period of relative stationarity coinciding with the decrease in wideband power (**Figure 6D, Supplementary Figure 9B**). To assess properties of oscillatory activity in these subjects, we detected spindle band oscillations. Such oscillations were present in all ages, consistent with delta brushes in the youngest subjects and sleep spindles in the older subjects (**Figure 6E**). We calculated spatial extent of these oscillations as the co-occurrence of spindle band activity across EEG electrodes, and found that this activity also shifted from highly localized to broader cortical expression around an inflection point between 42-47 weeks (**Figure 6F, Supplementary Figure 9C**). We also found evidence for significant cross frequency coupling of higher frequencies with the spindle band emerging after 47 weeks post-gestation (**Figure 6G-H;** n = 702 spindle band oscillations from 13 subjects; Kruskal Wallis Chi Square for AUC = 138.4; p < 0.001). Taken together, these data support the existence of a conserved transient quiescent period in development that dramatically shifts functional properties of neural networks.

**Figure 6:**
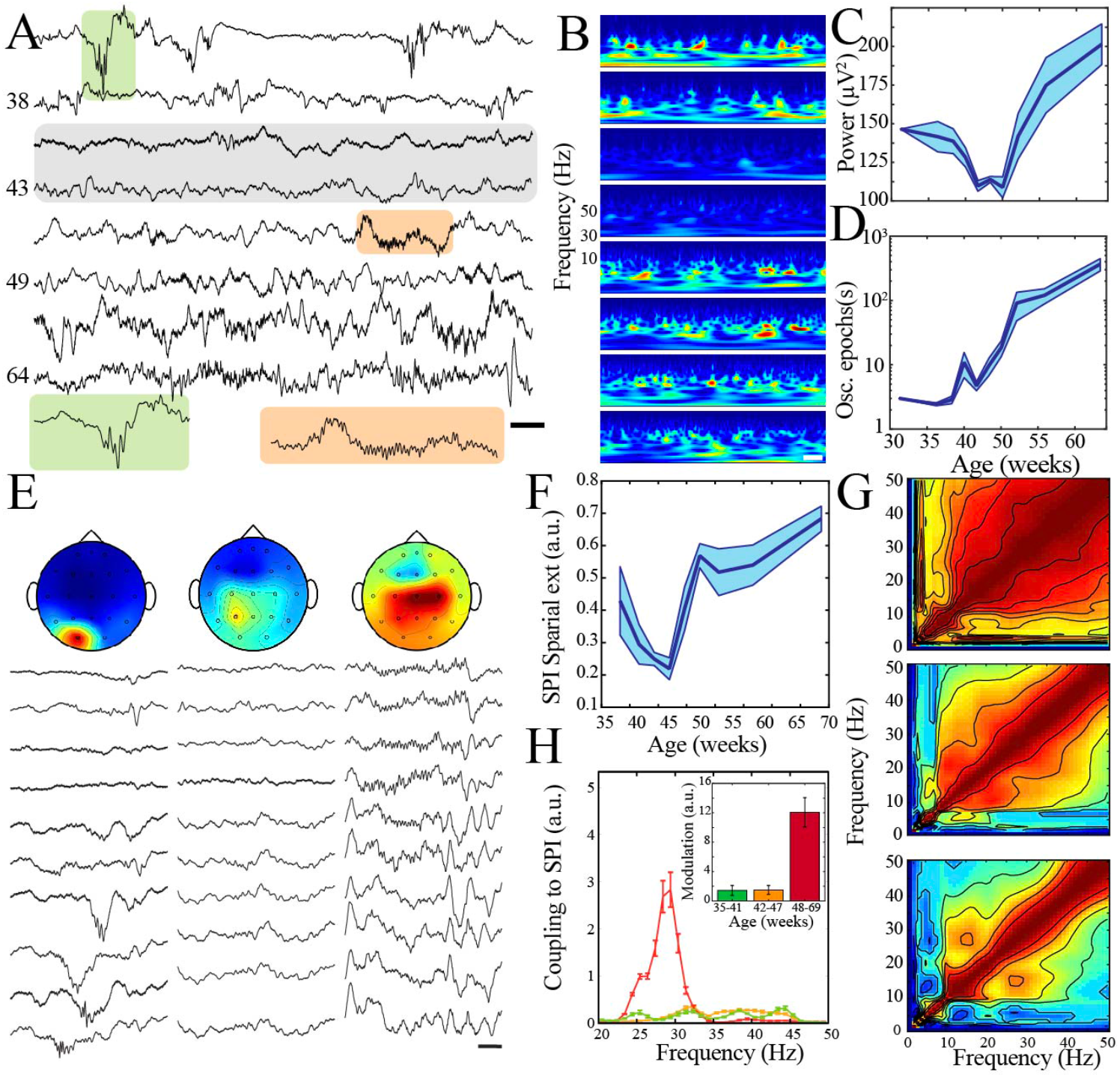
A transient quiescent state shifts network dynamics in humans. A) Sample raw traces from human subjects revealing a relative paucity of activity between 42-47 weeks during quiet/NREM sleep (gray shaded box). Orange shaded box is expanded to reveal characteristic appearance of infantile sleep spindles. Scale bar 1 s. B) Sample spectrograms from human subjects revealing a relative paucity of activity between 42-47 weeks during quiet/NREM sleep. Power scale was normalized across sessions. Scale bar 1 s; ages correspond to traces in A). C) Wideband (WB) power changes non-linearly across development (blue line = mean; shaded blue areas ± SE), with a local minimum between 42-47 weeks post-gestational age (n = 55 babies). Inset shows probability of a local minimum located at each age for datapoints resampled with replacement across days (grey) compared to datapoints resampled with replacement within days (orange), confirming existence of local minimum between 42-47 weeks. D) Duration of oscillatory epochs increases non-linearly across development (blue line = mean; shaded blue areas ± SE) with a local minimum between 42-47 weeks post-gestational age (n = 55 babies). Inset shows probability of a local minimum located at each age for datapoints resampled with replacement across days (grey) compared to datapoints resampled with replacement within days (orange), confirming existence of local minimum between 42-47 weeks. E) Sample raw traces of detected spindle band oscillations across subset of EEG electrodes from babies aged 37 weeks (left), 43 weeks (middle), and 58 weeks (right) post-gestational age. Scale bar 1 s. Head models show corresponding localization of spindle band power across EEG electrodes in conventional 10-20 placement. Power was normalized across sessions. F) Spindle band oscillation spatial extent changes non-linearly across development (blue line = mean; shaded blue areas ± SE), with a local minimum between 42-47 weeks post-gestational age (n = 55 babies). Inset shows probability of a local minimum located at each age for datapoints resampled with replacement across days (grey) compared to datapoints resampled with replacement within days (orange), confirming existence of local minimum between 42-47 weeks. G) Frequency comodulation at the time of spindle band oscillations from 36, 42, and 56 week post-gestational age subjects. H) A significant, discrete peak of cross-frequency coupling with spindle band oscillations emerges after 47 weeks (n = 702 spindle band oscillations from 13 subjects). Inset shows quantification of area under the curve for coupling to spindle band oscillations between 25-35 Hz.

## Discussion

We demonstrate that maturation of cortical network dynamics is enabled by a discrete, relatively quiescent, evolutionarily conserved transition period during early postnatal development. During this period, oscillations and neural spiking display a nadir that signals the shift from local, loosely correlated, prominently sensory-driven patterns to internally organized, spatially distributed, and temporally precise activity. Such a developmental inflection point may be necessary to enable expression of mature mechanisms for computation within cortical networks.

Self-organization in complex systems is often mediated by processes that can facilitate sudden transitions in system behavior. Our results provide evidence for such a transition in developing cortex of rodents and humans. Because rodent neurodevelopment is temporally compressed relative to humans, an inflection epoch that lasts 1-2 days in mouse pups could conceivably extend over weeks in humans (*33*). Prior to this inflection point, oscillatory activity was intermittent with non-uniform cycle-to-cycle waveforms and high broadband frequency coherence. Oscillatory activities after the transition became nearly uninterrupted, exhibiting coupling at discrete frequency bands and increased waveform symmetry. Based on these similarities in oscillatory patterns, we infer that the neural spiking features we observe through invasive recordings in mouse pups could also extend to normally developing human infants, where such recordings are not possible. Neural spiking was more efficiently recruited to oscillatory activity with consistent, strong phase-locking, and higher temporal precision. These features are characteristic of adult neural networks engaged in information processing (*16*). Thus, a transient period of developmental perturbation may be an evolutionarily conserved mechanism to facilitate a substantial shift in cortical network operations.

Our results suggest that the cortical network dynamics shift to permit temporally precise neural spiking after the transition period. Significantly synchronized firing within 10-50 ms intervals was absent prior to this period, and robustly emerged afterward. This result is consistent with observed increase in pair-wise correlations of spike activity in visual cortex over development (*48*). Although recruitment of neural spiking during epochs of spindle burst oscillations in immature rodents likely co-activates neurons that share similar afferent inputs (*7, 9–11*), more precise coordination is required for establishment of flexible, defined neural assemblies (*49*), and the capacity of circuits to express sequential activation of these assemblies is developmentally regulated (*50*).

We found that the neural population activity demonstrated correlated suppression over a progressively shorter duration (from approximately 500 to 200 ms) with maturation. This process is in keeping with the progressively decorrelated activity patterns observed when analysis is performed on the timescale of calcium indicators (*51, 52*). When oscillations occur in discrete frequency bands, the possibility for complex comodulation arises. This cross frequency coupling has been linked to information processing in the mature brain (*13–15*). We demonstrated that prior to the transient quiescent period, the dominant cortical oscillations (centered on spindle band) exhibit high comodulation extending from 1-100 Hz. During this period, broadband comodulation abruptly decreased and was subsequently replaced by discrete gamma oscillations. Therefore, the oscillatory infrastructure to support cross frequency coupling arises concomitantly with fine time-scale neural spiking synchronization and robust phase-locking of these spikes to LFP, suggesting an emergence of adult-like information processing capacity at this developmental timepoint.

Despite sharing a peak frequency, immature and mature spindle band oscillations are highly differentiable on the joint basis of waveform, power, and spatial extent. We found that although of high amplitude, spindle band frequency oscillations in the most immature mouse pups were highly and uniformly confined to localized regions of cortex (200-500 µm) roughly corresponding to cortical columns (*53, 54*). Similarly, human delta brushes exhibited restricted localization even given the limited spatial resolution of clinical EEG. During the transition period, spindle band oscillations in mouse pups had higher spatial extent, but markedly reduced amplitude, perhaps facilitated by increased intracortical synaptogenesis and maturation of corticothalamic afferents (*23*). The transition period in humans was marked by transiently decreased spatial extent of spindle band oscillations, which we hypothesize is related to attenuation of low power activity by the intervening tissue between cortical surface and scalp electrode. We observed that mature spindle band activity across species was characterized by increased and customizable spatial extent. Thus, the end of the transient quiescent period marks capacity for oscillatory frequencies that were previously confined to local processing units to extend into broader cortical areas with the potential to establish intercortical communication.

Functional inhibitory connections regulate physiologic oscillations in mature neural networks enabling submillisecond precision of spike timing and facilitating inter-regional synchronization (*55–57*). In contrast, interneurons do not substantially pace immature oscillations during the first postnatal week, though they can be recruited to and contribute to spatial properties of this activity (*7, 58*). Functional feedforward inhibition rapidly matures around the beginning of the second postnatal week in mice, resulting in emergence of adult-like hyperpolarizing synaptic potentials and precisely timed action potentials in glutamatergic neurons within the microcircuit (*24*). This timepoint, derived from *in vitro* data, temporally overlaps with the onset of adult-like neural network activity we observe *in vivo*. In the human brain, when mature functional inhibition arises is not known, but ongoing migration of cells that differentiate into interneurons in the months after birth (*59*) suggest the possibility of a postnatal transition to such network activity. However, given the multitude of anatomic and functional changes that occur during the early postnatal epoch, it is likely that the shift in network operating mode we observe is mechanistically multifactorial.

Neural processes in the adult brain are characterized by a specific constellation of network dynamics that are necessary for execution of complex behaviors and cognition (*60*). Our results suggest that the capacity for these processes simultaneously emerges abruptly after a distinct transition period characterized by relative neural quiescence. This transient quiescent period may be critical to shift the neural network operating mode from a predominantly external stimulus driven state to one that facilitates the internal representations associated with planning and executive function.

## Materials and Methods

### Probe fabrication

PEDOT:PSS (Clevios PH1000) was purchased from Heraeus. Ethylene glycol, (3-glycidyloxypropyl)trimethoxysilane (GOPS), 4-dodecyl benzene sulfonic acid (DBSA), 3-(trimethoxysilyl)propyl methacrylate (A-174 silane) were purchased from Sigma-Aldrich. Micro-90 concentrated cleaning solution was purchased from Special Coating Services. AZnLOF2020 (negative photoresist), AZ9260 (positive photoresist), AZ 400K and AZ 300MIF (metal ion free) developers were acquired from MicroChemicals, Merck. To create the PEDOT:PSS films, a mixture of aqueous dispersion (Clevios PH1000) and ethylene glycol was prepared and mixed with GOPS (1 wt%) and DBSA (0.1 wt%). The fabrication process involved deposition and patterning of parylene C, evaporation of Au for electrodes and interconnects and PEDOT:PSS films coating. Parylene C (1.2-µm-thick) was deposited on quartz wafers (100 mm outer diameter (O.D.), thickness of 1 mm) using an SCS Labcoater 2. For the metal lift-off, AZnLOF2020 negative photoresist was spin-coated at 3,000 r.p.m. on the substrate, baked at 105 °C for 90 s, exposed to ultraviolet light using a Suss MA6 Mask Aligner and developed with AZ 300MIF developer. Metallic layers (10 nm Ti, 150 nm Au) were deposited using an e-beam metal evaporator (Angstrom EvoVac Multi-Process) and patterned by soaking the substrate in a bath of resist remover. A second layer of parylene C (insulation layer) was deposited to a thickness of 1.2 µm and its adhesion to the bottom layer was enhanced by the addition of 3-(trimethoxysilyl)propyl methacrylate (A-174 silane) during chemical vapour deposition. An anti-adhesion agent (5 wt% Micro-90 diluted in deionized water) was deposited on the substrate followed by an additional sacrificial layer of parylene C that was subsequently used for the peel-off process. The stacked layers were patterned with a layer of AZ9260 positive photoresist and dry etched with a plasma-reactive ion-etching process (Oxford Plasmalab 80; 180 W, 50 sccm O2 and 2 sccm SF6) to shape electrode area and electrical contact pads. Specifically, AZ9260 was spin-coated at 5,000 r.p.m., baked at 115 °C for 90 s, exposed using a Suss MA6 Mask Aligner and developed with AZ400K developer (1:4 with deionized water). To obtain clean etching of large areas, the electrical contact pads were guarded by an extra layer of AZnLOF2020 (3,000 r.p.m.) between the metal layer and the silane-treated parylene. Each pad was etched along its perimeter and the parylene deposited on top of the photoresist was removed once immersed in acetone bath. Finally, PEDOT:PSS was spin coated on top of the electrodes and patterned by peeling off the last parylene layer. The probes had 119 electrodes regularly spaced on a diagonal square centered lattice with pitch of 77-177 µm and regularly interspersed perforations were created. Total device thickness was 4 µm.

### Animal Surgical Procedure

All animal experiments were approved by the Institutional Animal Care and Use Committee at Columbia University Irving Medical Center. Data from 108 Swiss-Webster mouse pups (3-9 g, 5-14 days of age) that underwent intracranial implantation was used for neurophysiological analysis. Pups were kept on a regular 12 h-12 h light-dark cycle and housed with mother before surgery. No prior experimentation had been performed on these mice. Anesthesia was induced by hypothermia in ice for P5-6 pups, and with 3% isoflurane for pups aged P7-14. Pups were then comfortably placed on a customized platform for head fixation, surgery, and recording. All animals were maintained under anesthesia with 0.75-1.5% isoflurane during surgery. Pre-and post-operative systemic and local analgesics were employed. Electrodes fabricated from fine gauge wire were inserted into the subcutaneous tissue of the neck and abdomen. These electrodes continuously registered electrocardiography (ECG) and electromyography (EMG) signals. A custom metal plate was attached to the skull to provide head fixation. For NeuroGrid implantations, a craniotomy was centered over primary somatosensory cortex and the array was placed over the dura. For silicon probe implantations, H32 NeuroNexus 32-site single-shank probes were used. A small cranial window over primary somatosensory cortex (approximately AP −1.5 mm, ML 2 mm; adjusted proportionally based on pup age) was opened, and the probe was stereotactically advanced until all electrodes were inserted below the pial surface (900 µm). Post-operatively, the pup was comfortably positioned, provided with familiar olfactory cues, and placed in a custom recording box with temperature and humidity control.

### Rodent neurophysiological signal acquisition

Neurophysiological signal acquisition was performed in the custom recording box, which was grounded to shield ambient electrical noise. Concurrent video monitoring was performed using a camera mounted inside the box. Pups recovered from anesthesia, as determined by increase of heart rate to plateau and onset of sleep-wake cycling, for at least 30 minutes before neurophysiological recordings were used for analysis. Neurophysiological signals were amplified, digitized continuously at 20 kHz using a local preamplifier (RHD2000 Intan technology), and stored for off-line analysis with 16-bit format. Spontaneous activity was acquired for 1-2 hours prior to euthanasia and perfusion.

### Perfusion and histology

At the end of experimentation, pups were euthanized with an overdose of pentobarbital (100 mg/kg). For the NeuroGrid experiments, the position of the array was visualized using light microscopy and the brain tissue underlying the corners of the array was stereotactically marked using a 30 gauge needle coated with chitosan compound (*31*). Pups were perfused with phosphate-buffered saline (PBS), followed by 4% paraformaldehyde. Brains were stored for 24–48 h in 4% PFA at 4°C prior to transfer to PBS solution. Cortices were flattened at room temperature (RT) after removal of the underlying midline structures and kept in 4% PFA for 24 h (4°C). Flattened cortices were sliced using a Leica vibratome vt1000S at 50 µm. The slices were then incubated for 1 h at RT in a blocking solution composed of PBS (0.01 M), 3% Normal Donkey Serum, and 0.3% Triton X-100. Slices were transferred to a solution containing anti-Vesicular Glutamate Transporter 2 (VGlut2) polyclonal guinea-pig antibody (AB2251-I Sigma Millipore) in a 1:2000 dilution and kept overnight at 4 °C. Subsequently, the tissue was washed and incubated with secondary antibody, Alexa Fluor 594 AffiniPure Donkey Anti-guinea pig IgG (H+L) (706-585-148) from Jackson Immunoresearch at 1:1000 dilution. Lastly, the tissue was incubated with DAPI (D9542-5MG; Sigma Aldrich) at 0.2 µg /mL and after washing, the brain slices were mounted with Fluoromount-G (00-4958-02, ThermoFisher). Fluorescent images were captured and tiled using a computer-assisted camera connected to an ECHO Revolve microscope. The images were adjusted for brightness and contrast and assembled into panels using ImageJ and Adobe Illustrator. The regularly spaced, hexagonal lattice of electrode spacing across the NeuroGrid enabled localization of each electrode relative to the marked corners of the array and primary sensory cortices. Localization of silicon probe electrodes to cortical region and layers was performed using a combination of intra-operative visualization and histological reconstruction. Probes were inserted under visual guidance, with the most superficial electrode placed just below the pia. Regular, precise spacing of electrodes along the shank enabled determination of each electrode’s depth from the cortical surface. Coronal brain slices stained with DAPI and vGlut2 permitted localization of probe implantation site relative to barrel cortex.

### Rodent neurophysiological signal pre-processing and state scoring

All recordings were visually inspected for quality, and channels that exhibited ongoing electrical or mechanical artifact were removed from analysis. Data was analyzed using MATLAB (MathWorks) and visualized using Neuroscope. Raw recordings were down-sampled to 1250 Hz to obtain local field potentials (LFP), and band-pass filtered between 250 Hz and 2500 Hz for analysis of neural spiking. Signals from EMG wires inserted into nuchal and ventral subcutaneous tissue were high-pass filtered at 300 Hz, rectified, and smoothed to obtain a power envelope. The mean of this high-pass filtered power was determined. Epochs consistent with high tone were identified when the signal surpassed a threshold defined by 1.5-3.5 standard deviations above this mean. Wakefulness was defined as a period of sustained high tone for at least 1.5 seconds. Myoclonic twitches involving contractions of the muscles of the head, forelimbs, hindlimbs, or trunk as observed on video recording were correlated with an increase in EMG signal that surpassed the threshold for < 1.5 seconds. Muscle atonia (consistent with sleep) was identified when the EMG signal remained below the threshold for at least 2 seconds. Sleep epochs were further classified as putative active or quiet sleep. Quiet sleep was identified when muscle atonia was paired with quiescence of behavior for at least 10 consecutive seconds. Active sleep was identified when muscle atonia was interspersed with myoclonic twitches. This approach has been shown to correlate with electrophysiologic markers of active and quiet sleep, and has been used for state-dependent developmental analyses (*20*).

### Human neurophysiological signal pre-preprocessing and state scoring

We retrospectively analyzed EEG recordings from 55 patients who underwent continuous monitoring with surface electroencephalography (EEG) as part of clinical diagnostic assessment. Analysis of these data were approved by the Institutional Review Board at Columbia University Irving Medical Center, and all data collection occurred at this institution. Each patient had EEG electrodes placed based on the internationally recognized 10-20 configuration. Electrode impedance was monitored and maintained within appropriate ranges as per American Clinical Neurophysiology Society Guidelines by certified clinical EEG technologists. Patients ranging in age from birth to 8 months who had at least 4 hours of high-quality continuous EEG monitoring were identified through the clinical electrophysiology database (Natus). Using the hospital admission record, patients were excluded if: i) the EEG was reported as abnormal by the reading electrophysiologist; ii) the patient had a known underlying neurologic or genetic condition; iii) the patient was diagnosed with a new neurologic condition during the course of the hospital admission; iv) corrected gestational age could not be determined from information contained in the medical record. Indication for EEG in eligible patients was most commonly paroxysmal movements concerning to parents for abnormal activity, with discharge diagnosis identifying normal movements/behaviors (e.g. sleep myoclonus, roving eye movements during REM sleep) or brief resolved unexplained events (BRUE; **Supplementary Figure 8**). Corrected gestational age, rounded to the nearest week, was used to classify patient age. If the phrase “full term” was employed in the medical record, corrected gestational age was calculated based on birth at 40 weeks. This approach allowed a precision of 2 weeks in corrected gestational age. Recordings always included both waking and sleep epochs. Three to five representative epochs of artifact-free quiet/NREM sleep were identified for subsequent analysis by a certified clinical electrophysiologist. Data was sampled at 256-512 Hz by clinical amplifiers. Raw data (referential) exported from Natus was used for all analyses. If electrical noise contaminated the recording, a 60 Hz notch filter was used. P3 or P4 electrodes, overlying the parietal lobe, were selected for spectral analysis. If these electrodes were non-functional, C3 or C4 electrodes, overlying the central sulcus, were used.

### Spectral analysis and oscillatory activity

To visualize spectrograms, a parametric autoregressive (order of one) whitening method was first applied to the data. For rodent data (sampled at 1250 Hz) and human data (sampled at 256-512 Hz), time-frequency decomposition was performed using analytic Gabor wavelet transform to enhance temporal resolution. Wavelet spectrograms centered at twitch times were normalized by z-scoring and averaged to obtain trigger-averaged spectrograms. Power was extracted for quantification from the magnitude of the analytic Gabor wavelet transform. Epochs of oscillatory activity were defined by presence of rectified signal amplitude above the 99^th^ percentile of the wideband noise floor.

### Modeling neural properties across developmental trajectories

In order to quantify the changing tendency of each feature over time, we modeled the trends with linear and polynomial regression models. To find the optimal fit that most accurately represented the overall trend without overfitting, we used leave one out cross validation to evaluate the models. The model with the minimum mean squared error was considered to have the best fit. When model fit predicted existence of local extrema, we used bootstrapping to estimate the location of these extrema for the trajectory of each property we investigated. For animal data, we resampled with replacement within each day from P5 to P14. For human data, we resampled within each 1-2 weeks. Age was jittered with a window of 0.5 days and 0.5 weeks for animal and human data respectively, to aid with fitting. We found the age corresponding to the first peak or trough of the regression line across development. This process was repeated 10000 times to generate a probability distribution of location of local extrema across the developmental trajectory. The same process was repeated using resampling with replacement across all timepoints of the trajectory (10000 iterations) to generate a null distribution for comparison.

### Spindle band oscillation detection

Spindle band oscillations were detected based on wavelet-derived power and duration parameters. Intervals of oscillatory activity containing spindle band power were first detected and a ratio of normalized autoregressive wavelet (Gabor)-based *P*_*AR*_ was then calculated to identify discrete spindle band oscillations. Oscillatory intervals were detected by comparing the spindle band power (*P*_*spi*_) of 8-25 Hz with threshold. Thresholds were determined relative to median absolute deviation 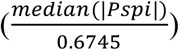 in the sleep intervals. Because this value varied markedly across development, thresholds were derived empirically but kept consistent across each age group. The value was decreased for silicon probe recordings in P5-7 pups compared to NeuroGrid recordings to maintain consistent detection despite quantifiably different signal to noise ratio. *P*_*AR*_ was then calculated based on the following equation:

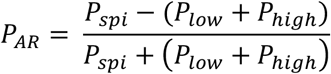

where spindle band power (*P*_*spi*_) was based on 8-25 Hz, low band power was based on 1-5 Hz (*P*_*low*_), and high band power was based on 30-80 Hz (*P*_*high*_). Spindle band events were identified when the ratio crossed −0.1 for a minimum of 300 ms and a maximum of 5 s and had a peak greater than zero. All detections were visually inspected for accuracy for each recording session and independently verified by an analyst blinded to detection parameters.

In human EEG data, spindle band oscillations were detected across all channels as in mice, using a constant threshold relative to median absolute deviation 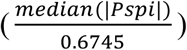 in the intervals of quiet/NREM sleep and *P*_*AR*_ was calculated as above. For subjects aged less than 40 weeks the low band power was based on 2-6 Hz to aid with accurate detection of delta brushes. The channel with maximal oscillation power in the spindle band was selected for further analysis.

### Spindle characterization

Spindle band oscillations were characterized by their power, spatial extent, and spindle asymmetry (*40*). Power was calculated by bandpass filtering each spindle band oscillation between 8-25 Hz. We then calculated the rectified value of Hilbert transformed filtered signals to derive the power envelopes. The median value of each spindle band oscillation was defined as its power. Spatial extent was calculated by determining the number of channels simultaneous expressing spindle band oscillations with a manually selected channel located in primary somatosensory cortex for mice and the channel with the highest spindle band power for humans (intersection of spindle start and end points > 300 ms and separation of spindle midpoints ≤ 50 ms on channels being compared). The spatial extent was expressed as a ratio of the number of channels with simultaneous occurrence and the total number of functional channels on the NeuroGrid. Data used for these calculations had a similar proportion and placement of channels located within primary somatosensory cortex. To quantify the sharpness of each spindle band oscillation’s peak and trough, they were first broadly bandpass filtered at 5-30 Hz. Rising and falling zero-crossing points were identified. Peaks were recognized as the maximum timepoint in raw data between a rising zero-crossing point and a falling zero-crossing point; troughs were recognized as the minimum timepoint in raw data between a rising zero-crossing point and a falling zero-crossing point. Sharpness of a peak was defined as the mean difference between the voltage at the oscillatory peak *V*_*peak*_ and the voltage at timepoints before (*V*_*before*_) and after the peak (*V*_*after*_), where the timepoint corresponded to a phase shift of approximately π (8 ms). Trough sharpness was defined similarly:

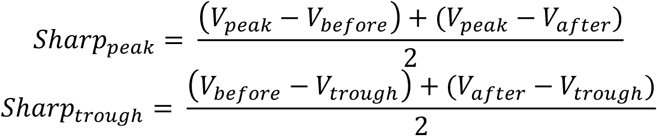

As the absolute difference between the extrema and the surrounding time points increases, the extrema sharpness increases. Extrema mean sharpness ratios (ESR) were calculated as a metric to quantify the sharpness asymmetry of each spindle. The ESR was defined in the following manner:

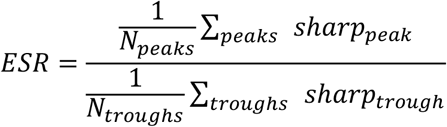

Larger deviation from 1 indicated greater sharpness asymmetry of the spindle.

### Spindle band activity clustering

In order to differentiate the spatiotemporal features of spindle band oscillations in immature (P5-7) and mature pups (P10-14), we applied a coarse tree model using spatial extent and ESR features. We trained the model with features extracted from 315 manually identified events from 12 pups. Five-fold cross validation was performed to avoid overfitting. The trained classifier was then applied to classify spindle band oscillations from different ages to test its ability to discriminate activities from immature and mature pups.

### Comodulogram and cross frequency coupling

Spectral analysis of the data was performed, and any data contaminated with electrical noise (60 Hz) was eliminated from analysis to prevent spurious correlations. Comodulograms of cross frequency coupling were computed based on a Gabor wavelet of raw data around the time of spindle band oscillation (4 s), followed by a Gaussian window convolution with duration matched to the frequency. The correlation matrix was calculated based on the magnitude of the wavelet components. Comparisons with p < 0.05 after Bonferroni-Holm correction for multiple comparisons were considered statistically significant. Columns of the correlation matrix corresponding to spindle band (10-20 Hz) were extracted and summated to obtain a curve representing the magnitude of frequency coupling to this band. Each local maximum coupling value was detected along with its peak prominence (how significant a peak is on account of its intrinsic value and location relative to other peaks) and peak width (frequency range with between half-prominence points). Peaks were thresholded based on prominence and width (Δf < 1/fc, peak prominence > 0.75). Wide-band coupling index was calculated by (peak value × peak prominence) / peak width. Area under curve (AUC) was calculated by integrated the summated coupling curves.

### Neural spiking analysis

Neural spiking was detected on each channel by thresholding negative peaks greater than 4 times the high pass filtered noise floor during epochs of sleep. Coincident spike times across a greater distance than expected for physiologic spike waveforms (∼ 300 µm) were presumed to be non-physiologic and eliminated from all channels. Electrodes were allocated based on histology and electrophysiologic characteristics into zones roughly corresponding to superficial (I-III) and deep (IV-VI) cortical layers. Electrodes from two non-contiguous sites (> 200 µm vertical separation) within the deep cortical layers were used for subsequent analysis. Because individual neuron action potentials could potentially be detected on more than one channel, spikes occurring on different channels within the same histologic zone < 2 ms apart were presumed to be duplicate detections and only the first spike was used for analysis. Inter-spike-intervals for the neural population were calculated as the absolute time between sorted spike times within the zone. A modified convolution method was used to determine 95% confidence intervals for population spike time autocorrelograms with a bin size of 10 ms. Peristimulus time histograms were calculated using a bin size of 50 ms with 95% confidence intervals determined from a shuffled distribution of spike times. To derive spike phase-locking to spindle band oscillations, the oscillatory epochs were first down-sampled to 125 Hz and narrowly bandpass filtered at 9-16 Hz. Phase was extracted using Hilbert transform. Phase bins π/24 were used for quantification of spike phase preference.

### Statistics

Statistical analysis was performed using a combination of open-source MATLAB toolboxes and custom MATLAB code. A modified convolution method was used to determine 95% confidence intervals for correlograms. Significance of phase-locking to spindle band oscillations was determined using alpha < 0.05 with a corresponding kappa value of > 0.1 Kuiper test (Circular Statistics Toolbox for MATLAB). Null distributions were generated based on 500 instances of data shuffling and used to calculate 95% confidence intervals. Probability distributions were compared using two-sample Kolmogorov-Smirnov tests with correction for multiple comparisons. Differences between groups were calculated using non-parametric rank sum (Wilcoxon) or ANOVA (Kruskal-Wallis with Bonferroni correction) depending on the nature of the data analyzed. When event size varied between groups, a random sample of events was selected from groups with larger event sizes to ensure group differences were not driven by degrees of freedom. Error bars represent standard error of mean. Significance level was *p* < 0.05.

## Acknowledgments

This work was supported by the Department of Neurology and Institute for Genomic Medicine at Columbia University Irving Medical Center as well as the School of Engineering and Applied Science at Columbia University. The device fabrication was performed at (1) Columbia Nano-Initiative, (2) Cornell NanoScale Facility (CNF), a member of the National Nanotechnology Coordinated Infrastructure (NNCI), which is supported by the National Science Foundation (Grant ECCS-1542081). SD received funding from the European Union’s Horizon 2020 research and innovation program under the Marie Skłodowska-Curie grant agreement No 799501. This work was supported by the NSF CAREER (1944415), CURE Taking Flight Award, Columbia School of Engineering as well as the Department of Neurology and Institute for Genomic Medicine at Columbia University Irving Medical Center. Thanks to the Churchland group for fruitful methodological discussion. We thank all Gelinas, Khodagholy, Buzsaki and Fishell laboratory members for their support.

All data needed to evaluate the conclusions in the paper are present in the paper and/or the Supplementary Materials. Additional data and code related to this paper may be requested from the authors

## Author Contributions

JNG, DK, SD, GB, and GF conceived the project. LM, SD, DK, JNG, GP and CM performed *in vivo* electrophysiological recordings and histological analysis. DK, CC, GS fabricated NeuroGrid probes. HY, LM, SD, JNG, DK analyzed electrophysiological recordings. JNG and DK wrote the manuscript with input from all authors.

## Declaration of Interests

The authors declare no competing interests.

---

**Supplementary Figure 1:**
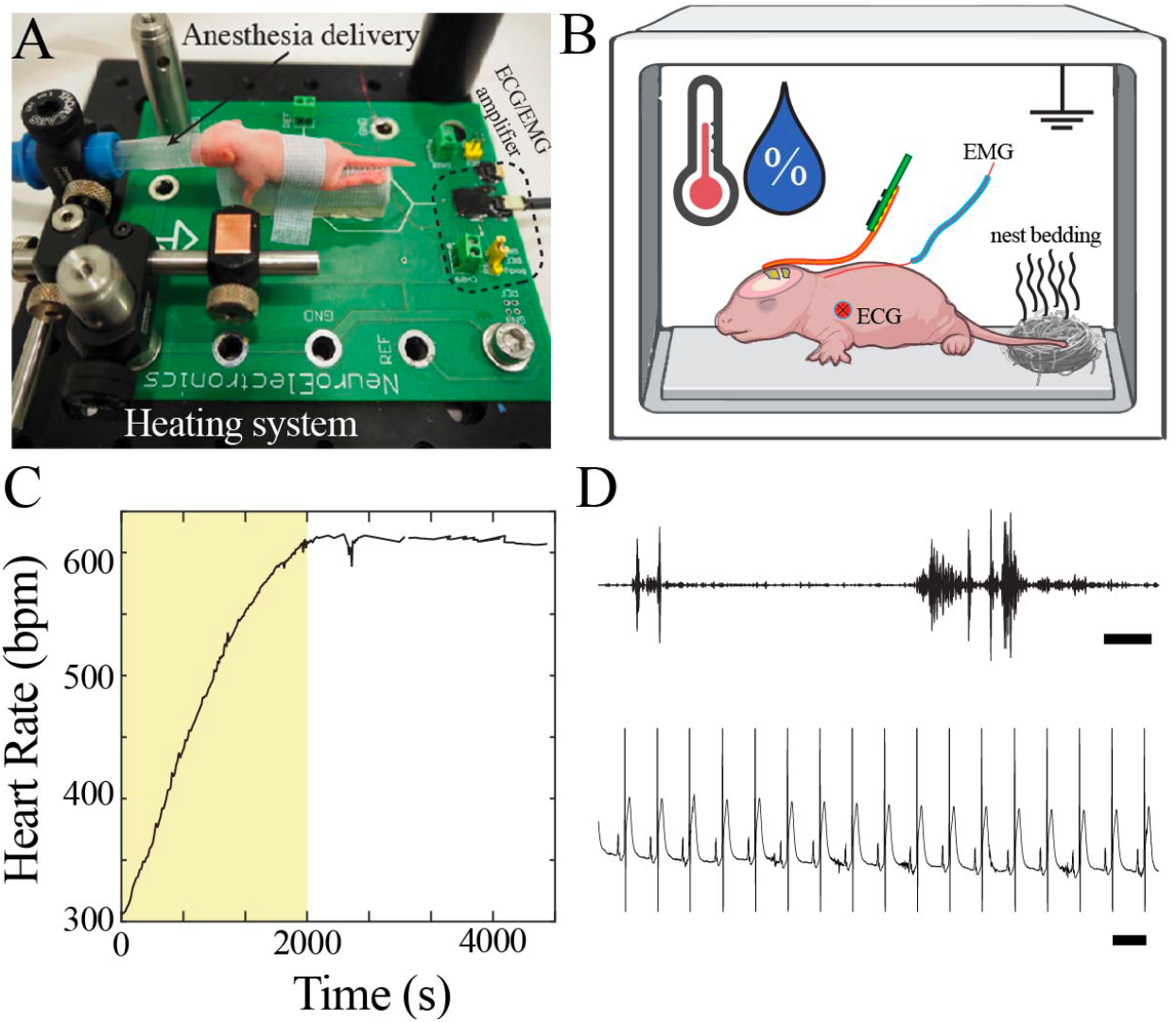
Neurophysiological recording in mouse pups. A) Customized surgical set-up with P7 pup for scale. B) Schematic of neurophysiological recording set-up with continuous recording of EMG and ECG, control of temperature and humidity, and placement of familiar olfactory cues. C) Heart rate increased to a plateau approximately 30 minutes after completion of surgical procedure. Data from epoch denoted by yellow box was not used for neurophysiological analysis. D) Sample high pass filtered trace from EMG electrode demonstrated interspersed bursts of EMG activity (upper, scale bar 1s). Sample raw trace from ECG electrode demonstrated regular heart rate (lower, scale bar 100 ms).

**Supplementary Figure 2:**
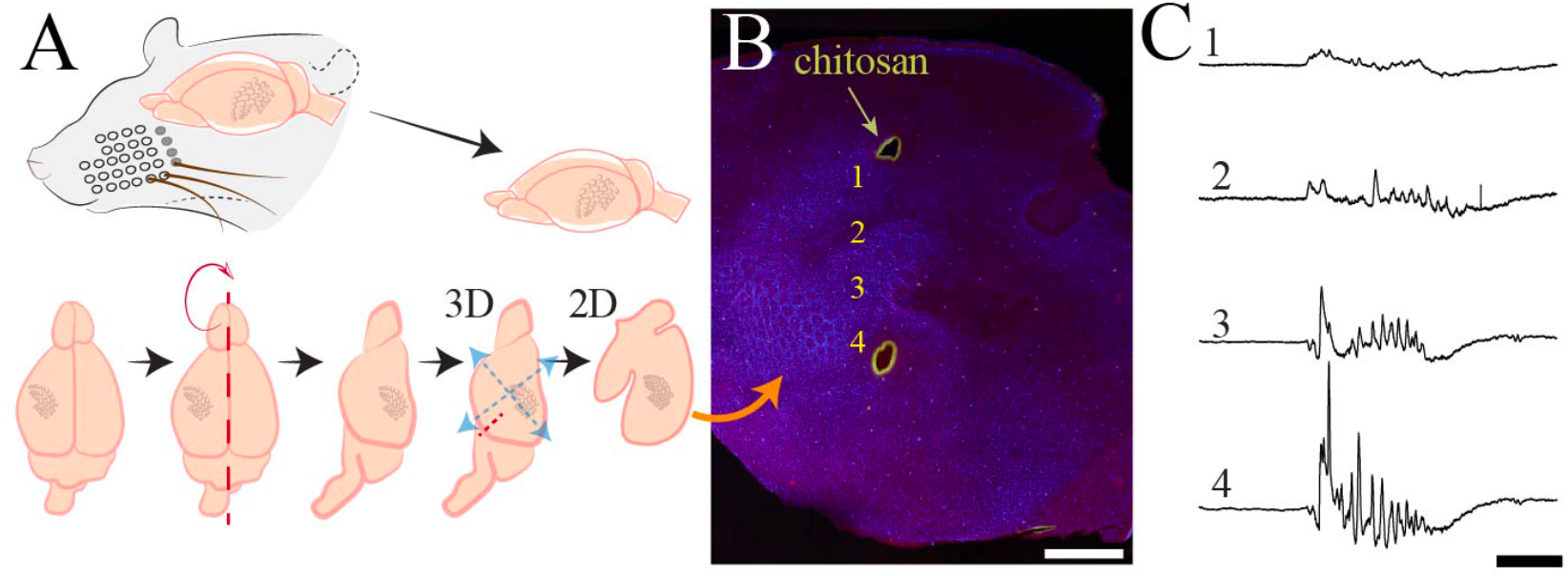
Histological processing and anatomical localization of NeuroGrid electrodes. A) Schematic demonstrating flattening of cortical mantle for immunohistochemical processing. B) Reconstruction of NeuroGrid location using chitosan marking (green) relative to somatosensory and visual cortices (visualized with vGlut2 + DAPI immunohistochemistry). Traces from numbered locations are represented in C). Scale bar 1.5 mm. C) Sample raw traces of a spindle burst oscillation from NeuroGrid electrodes at numbered locations in C). Scale bar 500 ms.

**Supplementary Figure 3:**
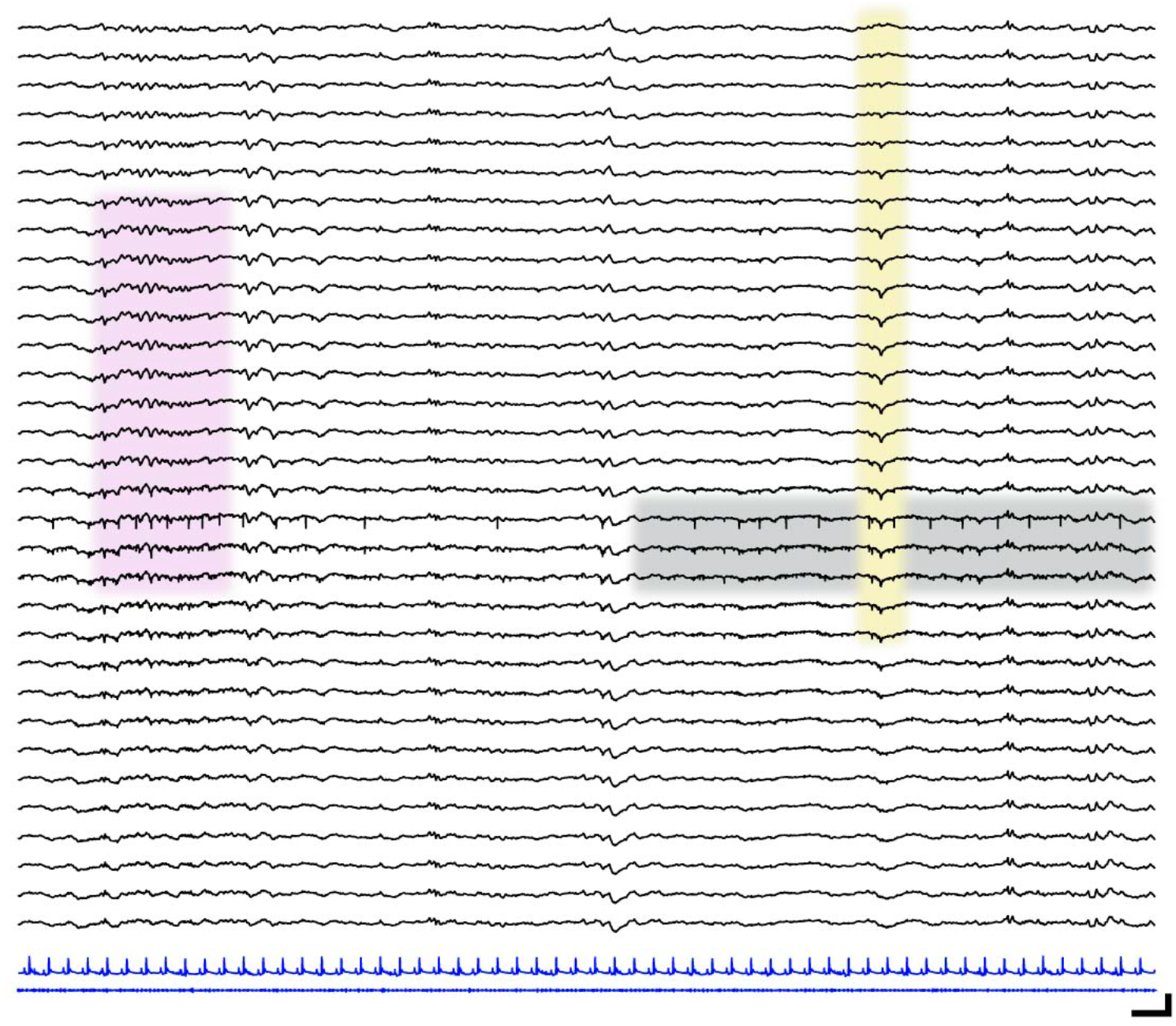
Localized neural spiking across cortical layers. Sample raw traces spanning cortical layers (black; upper = most superficial, lower = deepest) in a P13 mouse pup. Corresponding ECG (blue, upper) and EMG traces (blue, lower). Pink shaded box shows spindle band oscillation. Yellow shaded box highlights transcortical reversal of LFP waveform. Gray shaded box shows localized spiking activity. Scale bar 250 ms, 200 µV.

**Supplementary Figure 4:**
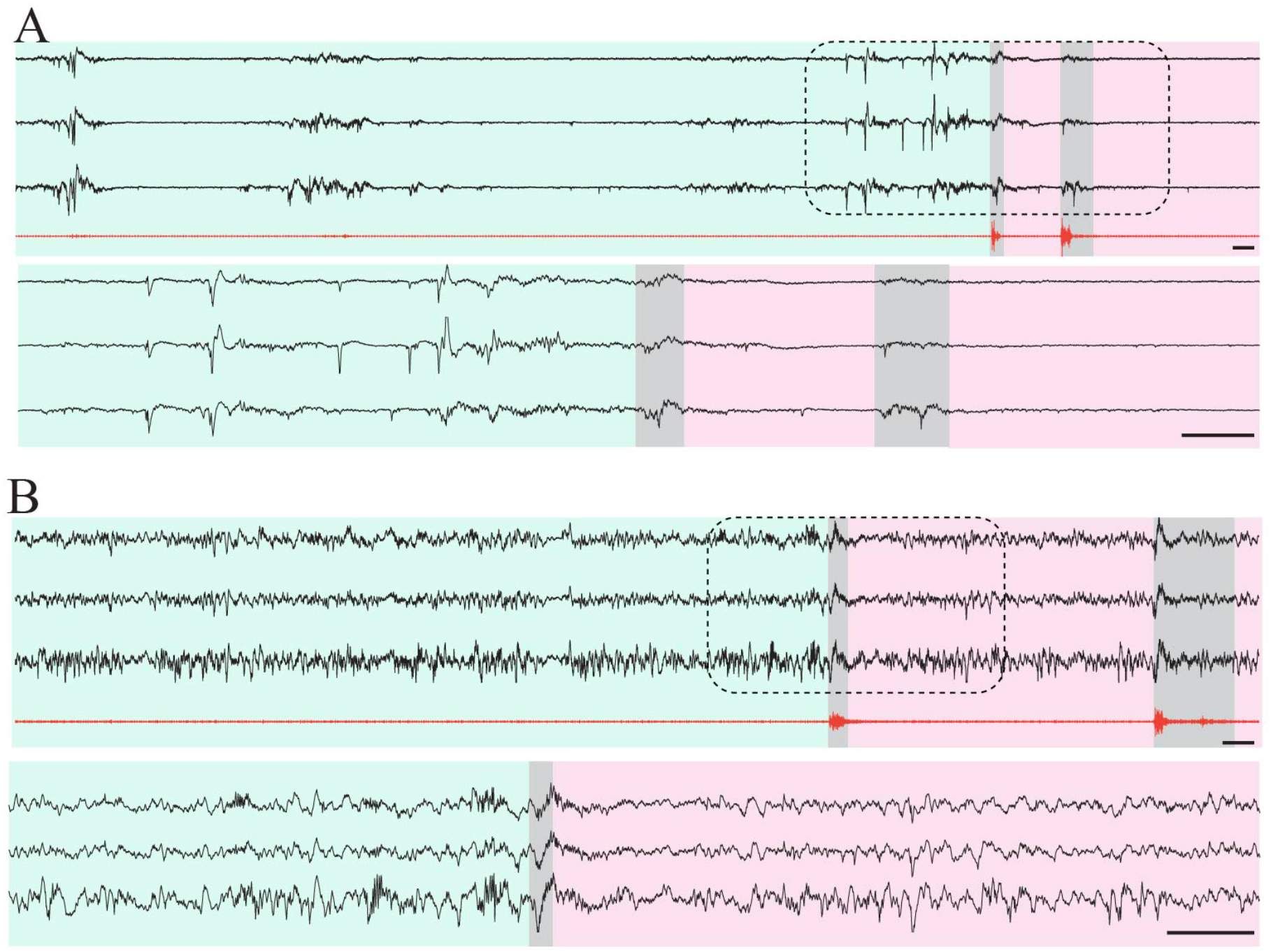
Isolation of putative quiet/NREM epochs in mouse pups. A) Three sample raw traces from P7 mouse pup (black) and high pass filtered EMG trace (red). Green shaded area indicates epoch of muscle atonia lasting > 10 seconds, consistent with putative quiet sleep. Scale bar 1 s. Grey shaded areas highlight myoclonic twitches. Pink shaded area indicates epoch with recurrent myoclonic twitches, consistent with putative active sleep. Lower three traces are zoomed in area denoted by dashed line box. Scale bar 1 s. B) Three sample raw traces from P14 mouse pup (black) and high pass filtered EMG trace (red). Green shaded area indicates epoch of muscle atonia lasting > 10 seconds, consistent with putative NREM sleep. Scale bar 1 s. Grey shaded areas highlight myoclonic twitches. Pink shaded area indicates epoch with recurrent myoclonic twitches, consistent with putative REM sleep. Lower three traces are zoomed in area denoted by dashed line box. Scale bar 1 s.

**Supplementary Figure 5:**
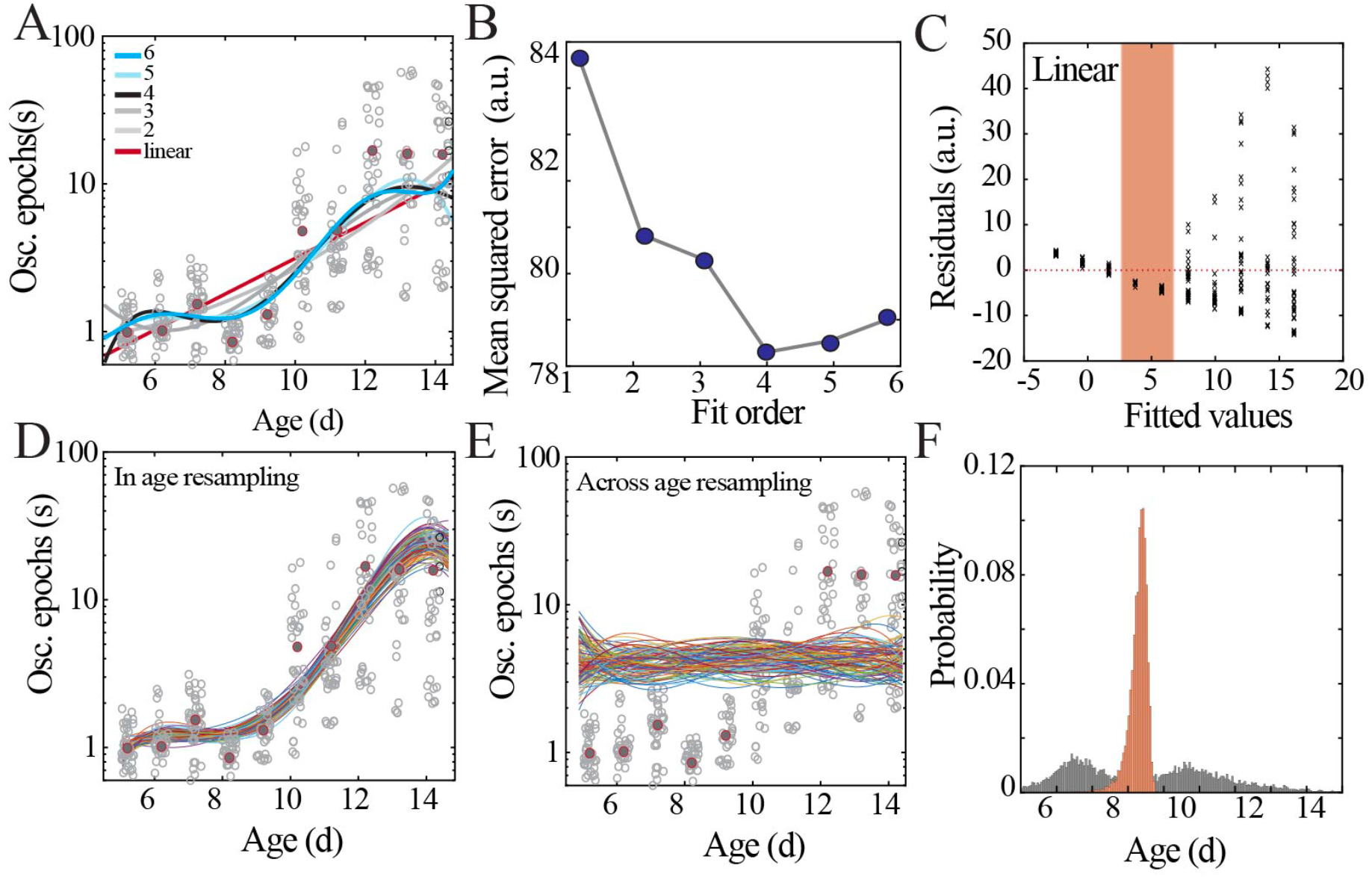
Model fitting and selection with bootstrapping to identify local minima. A) Data points from Figure 2C (gray circles) with superimposed regression models (linear, 2^nd^ to 6^th^ order polynomials). B) Leave-one-out cross-validation to evaluate fit of each model without over-fitting, quantified by mean squared error. C) Plot of residuals obtained from linear regression fitting; note systematic deviation from zero, especially at the beginning of the second postnatal week (orange shaded box). D) Distributions obtained by within age resampling with replacement (10000 iterations). For visualization purposes, only 100 randomly selected fits are displayed. E) Distributions obtained by across age resampling with replacement (10000 iterations). For visualization purposes, only 100 randomly selected fits are displayed. F) Comparison of nadir localization from distributions in D) and E) revealing significantly increased probability of local minimum between P8-9.

**Supplementary Figure 6:**
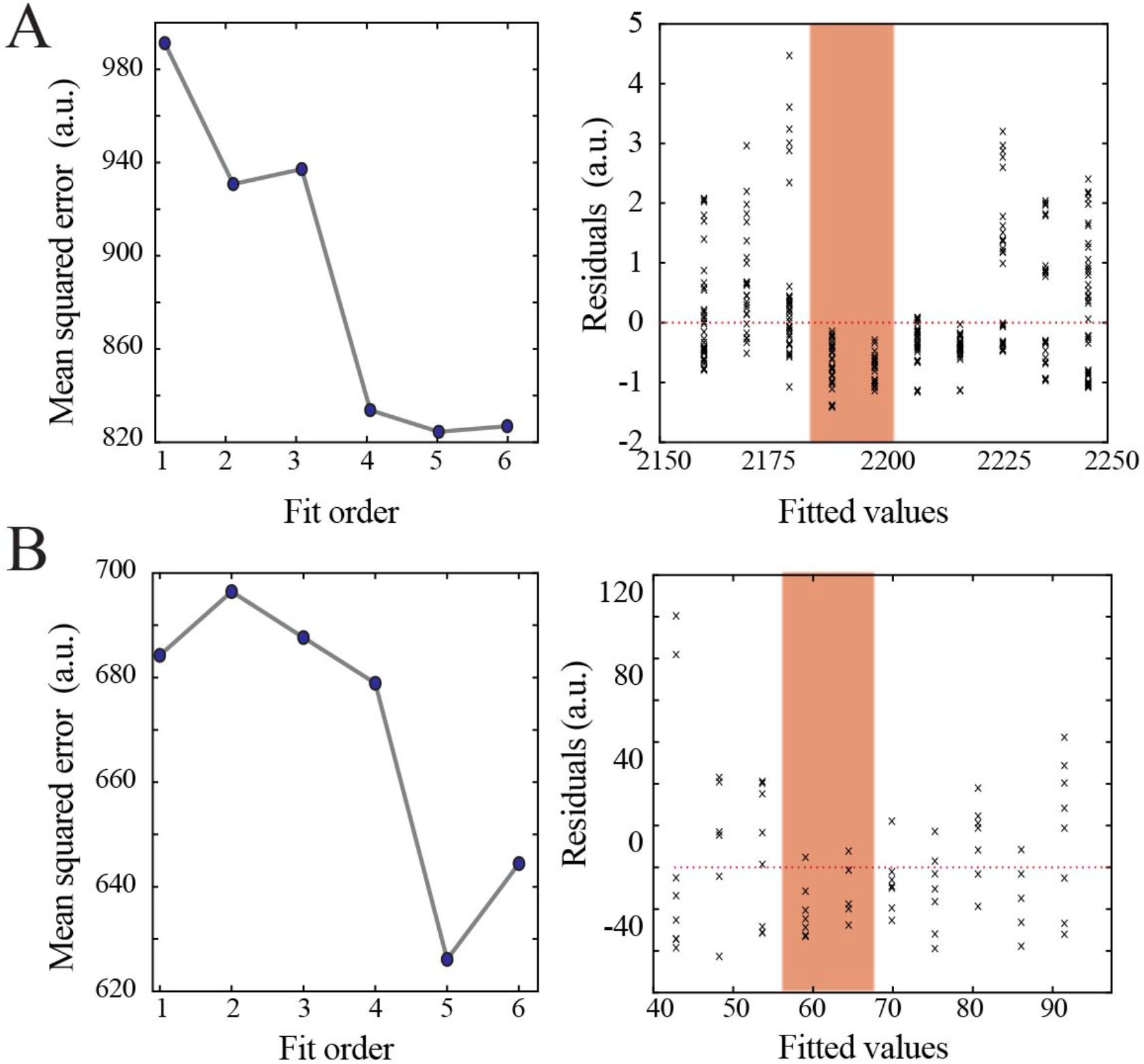
Model selection and verification of non-linear developmental trajectory for wideband power and spindle band oscillation properties in mice. A) Leave-one-out cross-validation to evaluate fit of each model for wideband power, quantified by mean squared error (left). Plot of residuals obtained from linear regression fitting (right); note systematic deviation from zero, especially at the beginning of the second postnatal week (orange shaded box). B) Leave-one-out cross-validation to evaluate fit of each model for spindle band oscillation properties, quantified by mean squared error (left). Plot of residuals obtained from linear regression fitting (right); note systematic deviation from zero, especially at the beginning of the second postnatal week (orange shaded box).

**Supplementary Figure 7:**
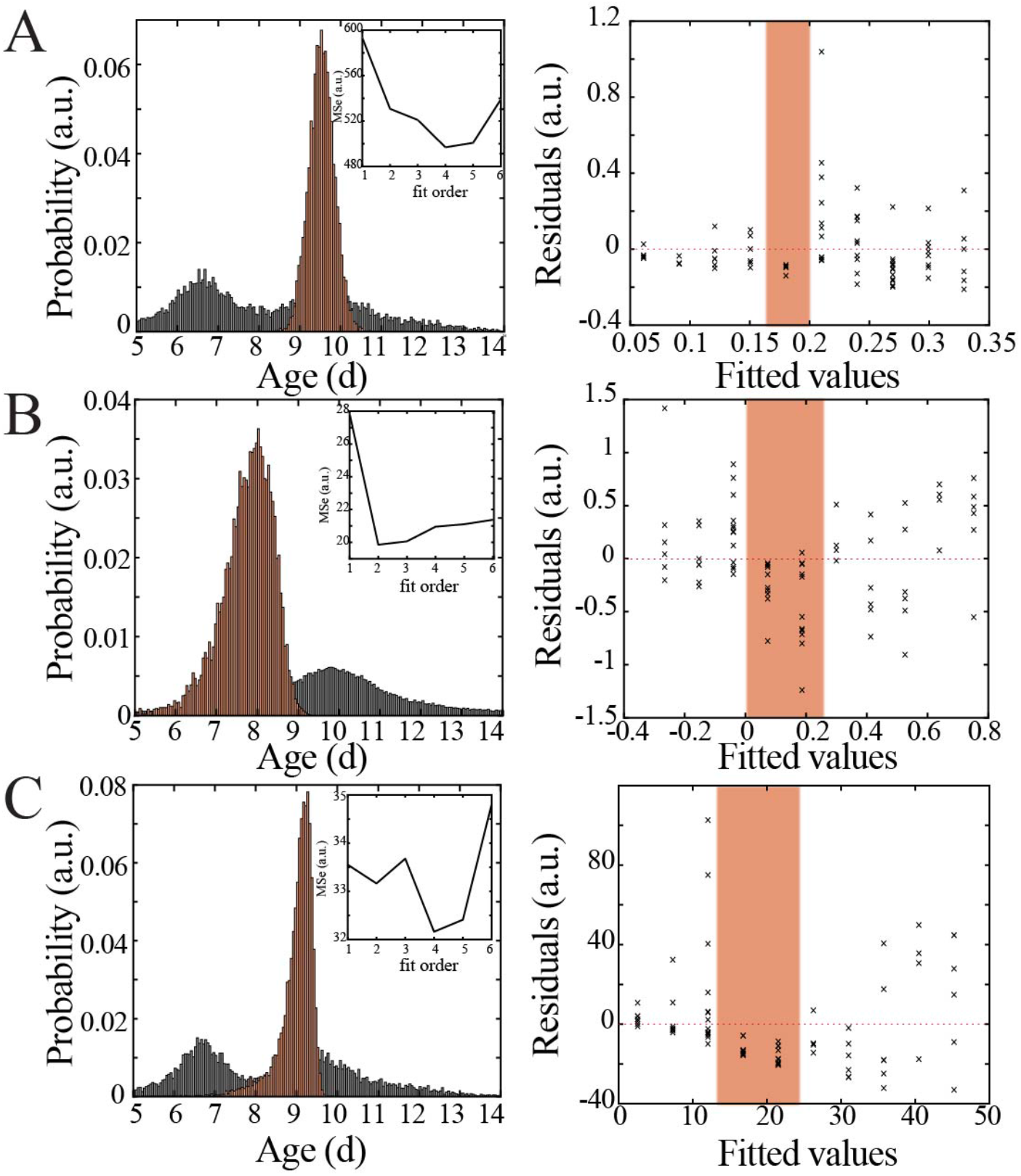
Bootstrapping, model selection and verification of non-linear developmental trajectory for spiking recruitment to spindle band oscillations, neural spiking rate, and interspike interval in mice. A) Comparison of nadir localization from distributions obtained by bootstrapping across ages (gray) and within ages (orange) revealing significantly increased probability of local minimum at the beginning of the second postnatal week (left) for spiking recruitment to spindle band oscillations. Inset shows leave-one-out cross-validation to evaluate fit of each model, quantified by mean squared error (MSe). Plot of residuals obtained from linear regression fitting (right); note systematic deviation from zero, especially at the beginning of the second postnatal week (orange shaded box). B) Comparison of nadir localization from distributions obtained by bootstrapping across ages (gray) and within ages (orange) revealing significantly increased probability of local minimum at the beginning of the second postnatal week (left) for neural spiking rate. Leave-one-out cross-validation to evaluate fit of each model, quantified by mean squared error (MSe). Plot of residuals obtained from linear regression fitting (right); note systematic deviation from zero, especially at the beginning of the second postnatal week (orange shaded box). C) Comparison of nadir localization from distributions obtained by bootstrapping across ages (gray) and within ages (orange) revealing significantly increased probability of local maximum at the beginning of the second postnatal week (left) for interspike interval. Leave-one-out cross-validation to evaluate fit of each model, quantified by mean squared error (MSe). Plot of residuals obtained from linear regression fitting (right); note systematic deviation from zero, especially at the beginning of the second postnatal week (orange shaded box).

**Supplementary Figure 8:**
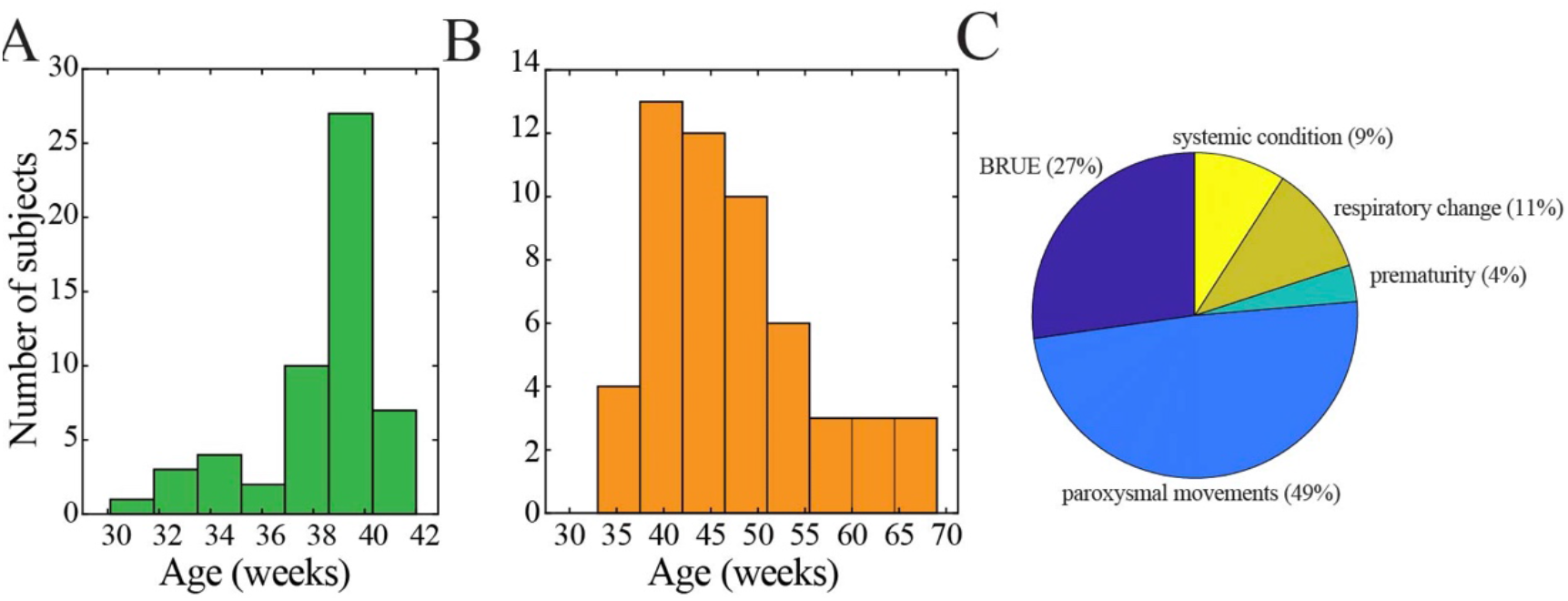
Demographic characteristics of human subjects. A) Histogram of age at birth. B) Histogram of age at time of continuous EEG monitoring. C) Clinical indication for continuous EEG monitoring (BRUE = brief resolved unexplained event).

**Supplementary Figure 9:**
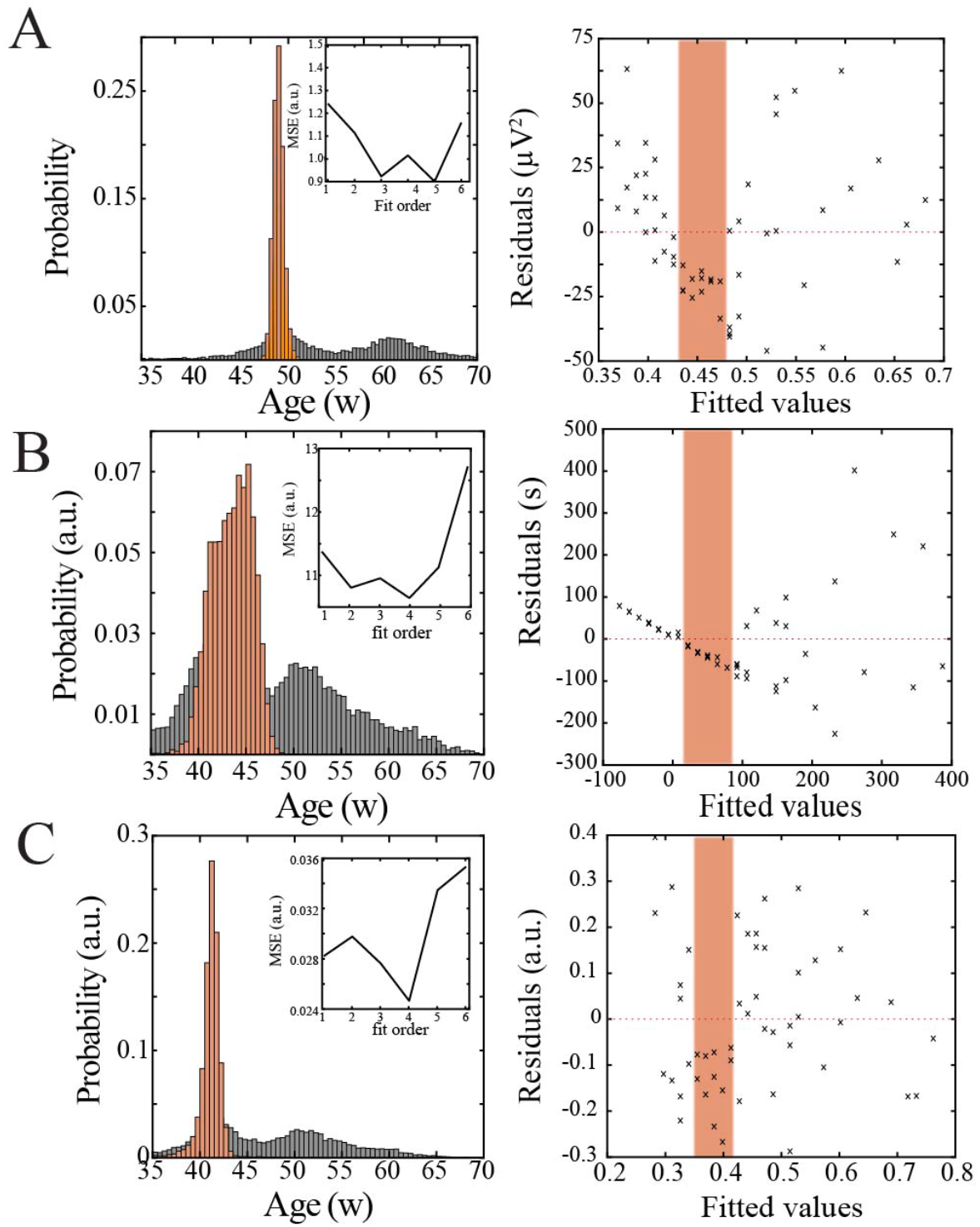
Bootstrapping, model selection and verification of non-linear developmental trajectory for continuity, wideband power, and spatial extent of spindle band oscillations in human subjects. A) Comparison of nadir localization from distributions obtained by bootstrapping across ages (gray) and within ages (orange) revealing significantly increased probability of local minimum between 42-47 weeks post-gestation (left) for wideband power. Inset shows leave-one-out cross-validation to evaluate fit of each model, quantified by mean squared error. Plot of residuals obtained from linear regression fitting (right); note systematic deviation from zero, especially between 42-47 weeks post-gestation (orange shaded box). B) Comparison of nadir localization from distributions obtained by bootstrapping across ages (gray) and within ages (orange) revealing significantly increased probability of local minimum between 42-47 weeks post-gestation (left) for continuity. Inset shows leave-one-out cross-validation to evaluate fit of each model, quantified by mean squared error. Plot of residuals obtained from linear regression fitting (right); note systematic deviation from zero, especially between 42-47 weeks post-gestation (orange shaded box). C) Comparison of nadir localization from distributions obtained by bootstrapping across ages (gray) and within ages (orange) revealing significantly increased probability of local minimum between 42-47 weeks post-gestation (left) for spatial extent of spindle band oscillations. Inset shows leave-one-out cross-validation to evaluate fit of each model, quantified by mean squared error. Plot of residuals obtained from linear regression fitting (right); note systematic deviation from zero, especially between 42-47 weeks post-gestation (orange shaded box).

## Notes

### Competing Interest Statement

The authors have declared no competing interest.

